# Deep Mutagenesis of a Transporter for Uptake of a Non-Native Substrate Identifies Conformationally Dynamic Regions

**DOI:** 10.1101/2021.04.19.440442

**Authors:** Heather J. Ellis, Matthew Chan, Balaji Selvam, Evan Walter, Christine A. Devlin, Steven K. Szymanski, L. Keith Henry, Diwakar Shukla, Erik Procko

**Author notes:** These authors contributed equally (Heather J. Ellis, Matthew Chan, Balaji Selvam).

## Abstract

The serotonin transporter, SERT, catalyzes serotonin reuptake at the synapse to terminate neurotransmission via an alternating access mechanism, and SERT inhibitors are the most widely prescribed antidepressants. Here, deep mutagenesis is used to determine the effects of nearly all amino acid substitutions on human SERT surface expression and transport of the fluorescent substrate APP+, identifying many mutations that enhance APP+ import. Comprehensive simulations of the entire ion-coupled import process reveal that while binding of the native substrate, serotonin, reduces free energy barriers between conformational states to promote SERT dynamics, the conformational free energy landscape in the presence of APP+ instead resembles Na^+^ bound-SERT, with a higher free energy barrier for transitioning to an inward-facing state. The deep mutational scan for SERT-catalyzed import of APP+ finds mutations that promote the necessary conformational changes that would otherwise be facilitated by the native substrate. Indeed, hundreds of gain-of-function mutations for APP+ import are found along the permeation pathway, most notably mutations that favor the formation of a solvent-exposed intracellular vestibule. The mutagenesis data support the simulated mechanism in which the neurotransmitter and a symported sodium share a common cytosolic exit pathway to achieve coupling. Furthermore, the mutational landscape for SERT surface expression, which likely filters out misfolded sequences, reveals that residues along the permeation pathway are mutationally tolerant, providing plausible evolutionary pathways for changes in transporter properties while maintaining folded structure.

## INTRODUCTION

The serotonin transporter (SERT/SLC6A4) catalyzes the reuptake of serotonin from the synapse to terminate synaptic transmission (*1*, *2*). As evidenced by the effects of pharmaceuticals and illicit drugs that inhibit SERT, and genetic studies of knockout mice (*3*) and natural human variants (*4–7*), dysregulation of serotonergic signaling is associated with broad psychopathological symptoms, including major depression, anxiety, obsessive compulsive disorder, post-traumatic stress disorder, and attention-deficit/hyperactivity disorder, amongst others (*8*). Similar to other closely related monoamine neurotransmitter transporters for dopamine (DAT/SLC6A3) (*9*, *10*) and norepinephrine (NET/SLC6A2) (*11*), SERT mediated serotonin (5-hydroxytryptamine; 5-HT) translocation is coupled to the favorable dissipation of a Na^+^ electrochemical gradient. While the stoichiometry for coupled transport remains unclear (*12*), it most likely involves the Cl^-^-dependent import of one Na^+^ and one positively charged 5-HT into the cell by SERT (*13–15*) using an alternating access mechanism (*16*, *17*), in which the transporter in an outward-facing (OF) conformation captures extracellular substrate, transitions through an occluded (OC) state where the central substrate binding sites become closed and inaccessible to the outside, and then an intracellular exit pathway opens in an inward-facing (IF) conformation for cytosolic release (*18*, *19*). SERT then returns to an outward-facing conformation through the antiport of one K^+^, together with possible movement of a proton, for an overall electroneutral cycle (*12*, *13*). Alternative stoichiometries and even channel-like conduction states may occur under different conditions (*15*, *20–23*).

Monoamine transporters and other neurotransmitter:sodium symporters (NSS) share a common architecture (*24*) known as the LeuT fold, named after the bacterial LeuT transporter, which was the first member structurally characterized at atomic resolution (*25*). Static crystal structures of human SERT and *Drosophila* DAT have revealed atomic details of OF and OC conformations bound to drugs and substrates (*26–31*). These structures have resolved two Na^+^ sites (Na1 and Na2) and the location of Cl^-^ binding, as well as specific contacts to the neurotransmitter aromatic ring and amine moieties within subsites (*32*) of the central orthosteric (or S1) binding site. In addition, structural, biochemical and pharmacological studies have shown that substrates entering the outward-facing extracellular vestibule engage an allosteric (or S2) site immediately adjacent to critical gating residues, before moving past the open gate to the orthosteric site (*26*, *33–35*). Cryo-EM analysis of detergent-solubilized SERT in the presence of ibogaine, a psychoactive plant product that stabilizes IF conformations, allowed for the characterization of OF, OC and IF-like states at moderate resolution (*19*), showing that regions already implicated in bacterial and mammalian NSS homologues through multiple methods (*18*, *36–41*) move to open a solvent-accessible intracellular vestibule. In particular, transmembrane helix (TM) 1a moves away from the helical bundle. These motions are supported by comprehensive HDX-MS analysis of LeuT (*42*) and SERT (*43*). However, the overall molecular mechanism for ion-coupled neurotransmitter transport remains incompletely understood. Key questions that remain unresolved are how stoichiometry is achieved and how does the neurotransmitter move from the orthosteric site to the cytosol. Cryo-EM structures of ibogaine-bound SERT only confirmed the general region for neurotransmitter exit, and the authors were uncertain as to the exact exit pathway amongst multiple possibilities (*19*). Understanding the full mechanism of transport in atomic detail will assist in understanding how genetic variants are associated with disease and may aid the development of drugs targeting discrete conformational states.

We recently described molecular dynamics (MD) simulations of the entire SERT- catalyzed import process of 5-HT (*44*), accomplished using adaptive sampling to efficiently explore conformational space. The calculated highest flux pathway for substrate binding events and conformational changes lead to several important conclusions, including the identification of key residues involved in substrate recognition at the extracellular vestibule and the prediction that the neurotransmitter exits along a pathway beginning at the Na2 site that is surrounded by TM1a, TM5, TM6b, and TM8.

Here, we use deep mutational scanning (*45*) to define the mutational landscape of SERT for surface expression, which indicates how mutations effect folding and escape from intracellular protein quality control (*46*), and substrate transport. Deep mutagenesis involves (i) creating a library of protein variants, (ii) using *in vitro* selection to enrich for the variants with high activity, and (iii) measuring the enrichment or depletion of variants in the mutagenesis library using next generation sequencing. Deep mutational scanning has proven insightful for indirectly interrogating protein conformational states in living cells (*46–49*), and datasets that include higher order mutations can even assist in modeling small protein structures with atomic resolution (*50*, *51*). We build on this body of work by now applying deep mutagenesis to a human transporter that undergoes substantial conformational change. By using a non-native substrate that is unable to facilitate isomerization of SERT to an inwards-facing conformation when compared with the native substrate, we find extensive mutational information that provide insight on dynamic regions in close agreement with simulation.

## RESULTS

### Mutational landscapes of SERT for plasma membrane localization and substrate import

We sought to understand how SERT sequence relates to conformational dynamics and mechanism using deep mutagenesis, which demands an effective selection strategy to distinguish low and high activity SERT sequence variants. This is not possible using transport assays with radioactive 5-HT. Instead, *in vitro* selection was based on the uptake of the fluorescent substrate and monoamine neurotransmitter analogue APP+ (4-(4-dimethylamino)phenyl-1-methylpyridinium) (Figure S1), permitting cells expressing active and inactive SERT variants to be readily distinguished and separated by fluorescence. We used a fluorescence-based transport assay to assess SERT activity by flow cytometry, which could then be seamlessly integrated with fluorescence-activated cell sorting (FACS) to screen libraries of SERT variants. To detect surface localized transporter, a synthetic gene encoding human SERT with a c-myc epitope tag (flanked by gly/ser-rich linkers) inserted between residues Asp216 and Asn217 of EL2 was constructed; the homologous site in *Drosophila* DAT features a large domain insertion. When expressed from an episomal plasmid in Expi293F cells (a suspension culture derivative of human HEK293 that has advantages for FACS-based selections of large libraries), surface localized transporter was detected with a fluorescent anti-myc antibody, while uptake of APP+ was simultaneously observed (Figures S2A and S2B). Validation of our construct showed that myc-tagged SERT has ion dependence consistent with untagged human SERT (*14*, *52–54*) (Figure 7E), appropriate neurotransmitter selectivity (Figure 7D), and exhibits a similar K_M_ for APP+ as for the endogenous substrate 5-HT (Figure 7E and Table 1). However, APP+ and 5-HT transport velocity of myc-tagged SERT are reduced 4-fold and 2-fold compared to the native protein, respectively, while insertion of a shorter gly/ser-rich sequence at the same site had no significant effect (Figure S2C and Table 1). We speculate that the bulkier myc tag with a longer gly/ser-rich linker (a 24 residue insertion in total) may partially obstruct substrate access to the extracellular vestibule, a limitation that we considered offset by the capability of measuring transport and surface localization simultaneously. Furthermore, a subset of mutations displaying enhanced APP+ transport discovered through deep mutagenesis were confirmed to also enhance activity in native untagged SERT (Figure S2D), and the presence of the myc tag is therefore very unlikely to have impacted scientific conclusions.

**Table 1.** 5-HT transport: K_M_ and V_max_ of SERT and mutants.

| Transporter | $K_M$ ($\mu\text{M}$)<sup>1</sup> | $V_{\max}$ (amol cell<sup>-1</sup> min<sup>-1</sup>) | n |
| --- | --- | --- | --- |
| native SERT | $2.1 \pm 0.7^{**}$ | $0.66 \pm 0.15$ | 3 |
| opt myc-SERT<sup>2</sup> | $1.3 \pm 0.6$ | $0.48 \pm 0.15$ | 4 |
| R79V | $0.25 \pm 0.07^*$ | $0.05 \pm 0.01^{**}$ | 3 |
| F88H | $0.14 \pm 0.05^*$ | $0.07 \pm 0.04^*$ | 3 |
| N173G | $1.2 \pm 0.5$ | $0.38 \pm 0.13$ | 4 |
| L406I | $1.1 \pm 0.5$ | $0.23 \pm 0.01$ | 4 |

<sup>1</sup> Kinetic parameters are mean ± SE. Data were evaluated using a 1-way ANOVA with post-hoc Tukey's comparison.
<sup>2</sup> Mutations are introduced into codon-optimized SERT with a c-myc tag in EL2.

Tagged SERT was mutagenized to generate nearly all single amino acid substitutions. To increase sampling during FACS-based selection, substitutions were split across two libraries spanning the N-(residues 2-325) and C- (residues 326-630) termini, each library containing ∼6,000 mutants. The libraries were diluted with carrier DNA encoding a viral factor for enhanced replication of the episomal plasmid (*46*) and transfected into Expi293F cells. A large excess of carrier DNA ensures that typically only one SERT coding variant is expressed in any cell (*48*), providing a tight link between genotype and phenotype. Cell libraries were incubated with APP+ at a concentration below the K_M_ and stained with anti-myc antibody conjugated to an orthogonal fluorochrome to simultaneously assess surface expression. Cells expressing SERT at the plasma membrane, or expressing SERT and transporting the fluorescent substrate, were collected via FACS (Figures S2E and S2F). After deep sequencing, mutation enrichment ratios were calculated by comparing variant frequencies from RNA transcripts in the sorted cells to the naive DNA library. These data experimentally define the mutational landscapes for SERT surface expression and APP+ transport (Figure 1).

**Figure 1.**
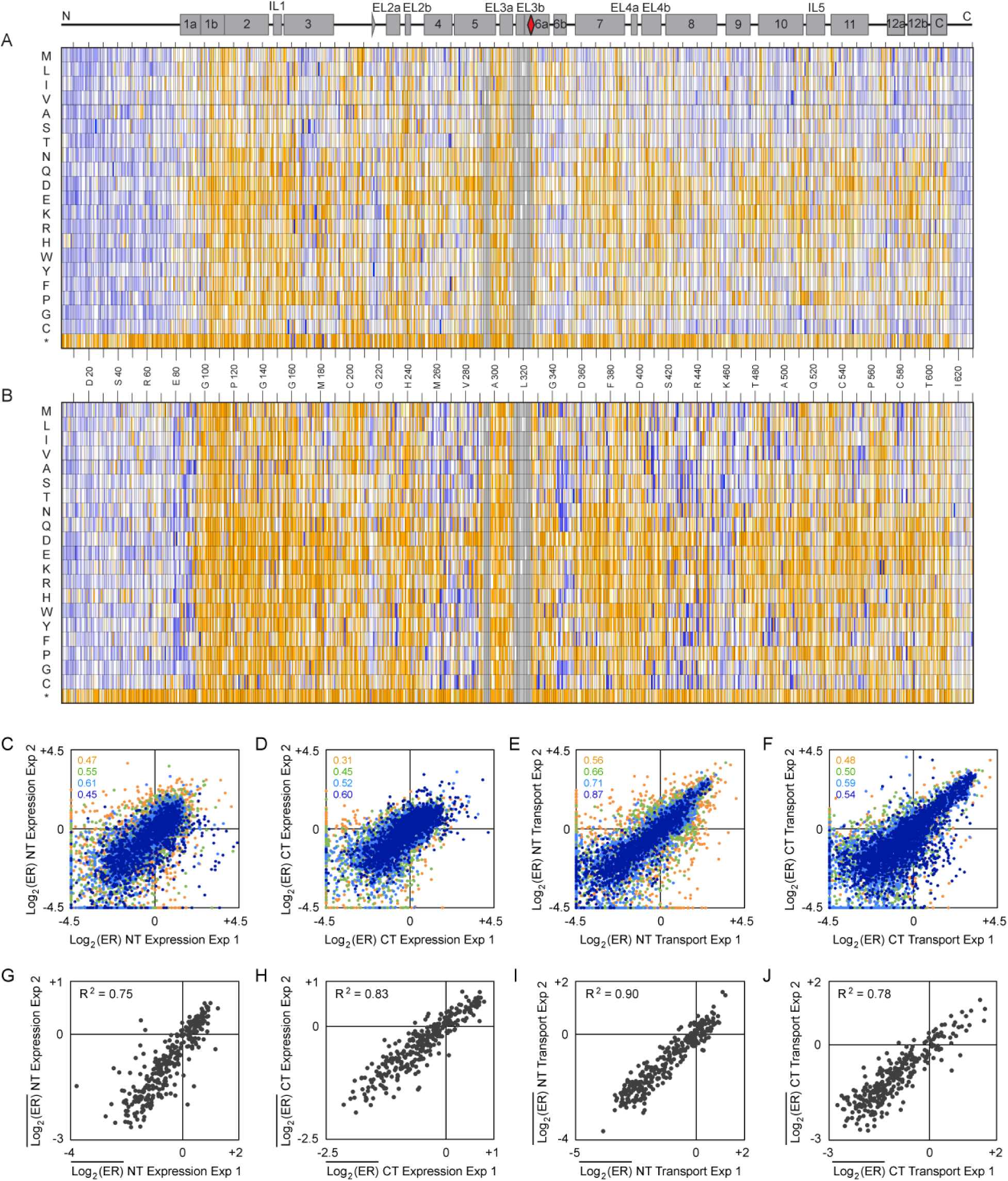
SERT mutational landscapes for surface expression and APP+ transport. **(A, B)** SSM libraries of SERT spanning N- and C-terminal residues (the red diamond in the secondary structure schematic at top indicates the dividing point between the libraries and the grey triangle at positions 216-217 indicates the placement of the myc tag) were sorted for (A) surface expression and (B) cell uptake of APP+. Log_2_ enrichment ratios are plotted from ≤ −3 (depleted mutations, orange) to 0 (neutral, white) to ≥ +4 (enriched, dark blue). Data are averaged from two independent experiments. Missing mutations (< 10 reads in the naïve libraries) are grey. *, stop codons. **(C-F)** Agreement between log_2_ enrichment ratios from replicated selections for surface expression of the SERT (C) N-terminal and (D) C-terminal SSM libraries, or for APP+ transport of SERT (E) N-terminal and (F) C-terminal SSM libraries. Agreement between independent experiments generally increases for more frequent sequence variants (orange, 10-99 reads in the naïve library; green, 100-199 reads; light blue, 200-399 reads; dark blue, 400+ reads). R^2^ values are correspondingly colored in the upper-left corners. **(G-J)** Agreement between conservation scores (defined as the mean of the log_2_ enrichment ratios at a single residue position) from replicated selections for surface expression of SERT (G) N-terminal and (H) C-terminal SSM libraries, or APP+ transport of (I) N-terminal and (J) C-terminal libraries.

Enrichment ratios are correlated between independently replicated selection experiments (Figures 1C-1F). Agreement between replicate experiments tends to increase for highly abundant sequence variants, presumably due to increased sampling. Metrics for mutational tolerance such as sequence Shannon entropy do not distinguish between positions that tolerate many neutral mutations versus positions that are hot spots for gain-of-function mutations. We therefore calculate a simple, unambiguous average of the log_2_ enrichment ratios for all 20 amino acid possibilities at each position, which we call a residue conservation score. Conservation scores are highly correlated between replicates (Figures 1G-1J) and define regions of sequence that are tightly conserved for function, or alternatively are under positive selection to change.

Mutational scans of transmembrane proteins for plasma membrane localization indicate whether mutations adversely impact folding and structural stability, as folded proteins are anticipated to preferentially escape intracellular quality control machinery (*46*). In the mutational landscapes for both SERT surface expression and APP+ transport, mutations within secondary structural elements are generally deleterious, especially polar mutations within transmembrane segments (Figures 1A and 1B). In mutational scans of G protein-coupled receptors (*48*), proline substitutions, which are disruptive to helical conformations, were depleted in regions (including termini) that have ordered structure, yet no such depletion of proline substitutions is observed in the cytosolic termini of SERT. Instead, the SERT cytosolic tails (N-terminal residues 1-75 and C-terminal residues 614-630 beyond the C-terminal helix immediately following TM12) tolerate most mutations, strongly indicating that these regions lack folded structure based on expression and substrate import alone, despite previous modeling to the contrary (*55*). However, we cannot exclude that these regions may have folded structure pertinent to regulatory mechanisms, such as structural elements induced by protein-protein interactions or post-translational modifications, that do not participate in our cell-based selection. In comparison, mapping conservation scores for each residue position to the crystal structure of SERT (*26*, *27*) highlights regions of folded structure under sequence constraints (Figure 2). Residues considered important for SERT trafficking, namely disulfide bonded cysteines (*56*) and a putative cholesterol interaction site (*57*), are moderately to highly conserved for high surface expression, whereas N-glycosylation site N208 is partially conserved and N217 is not (Figure S3). Moreover, co-evolved residues also tend to be well conserved in the mutational scan for surface expression, highlighting evolutionary constraints to maintain proper transporter folding and stability (Figure S4). The selection of SERT mutants for surface expression reveals conservation within packed hydrophobic cores ‘above’ and ‘below’ the central orthosteric binding site (Figure 2B), while the protein sequence along the permeation pathway is tolerant of many substitutions. This suggests mutations in the permeation pathway, which may be relevant in the evolution of new transporter specificities and coupled ion stoichiometries, may frequently occur without major disruption of the basic protein architecture and fold. Finally, contrary to predicted free energy values for membrane helix insertion (*58*), transmembrane helices surrounding the periphery of the transporter are only moderately conserved for expression, while helices in the core are highly conserved (Figure S5). The effects of mutations on SERT surface expression are therefore most closely related to the protein’s folded structure and not to the bioenergetics for membrane insertion of individual helices.

**Figure 2.**
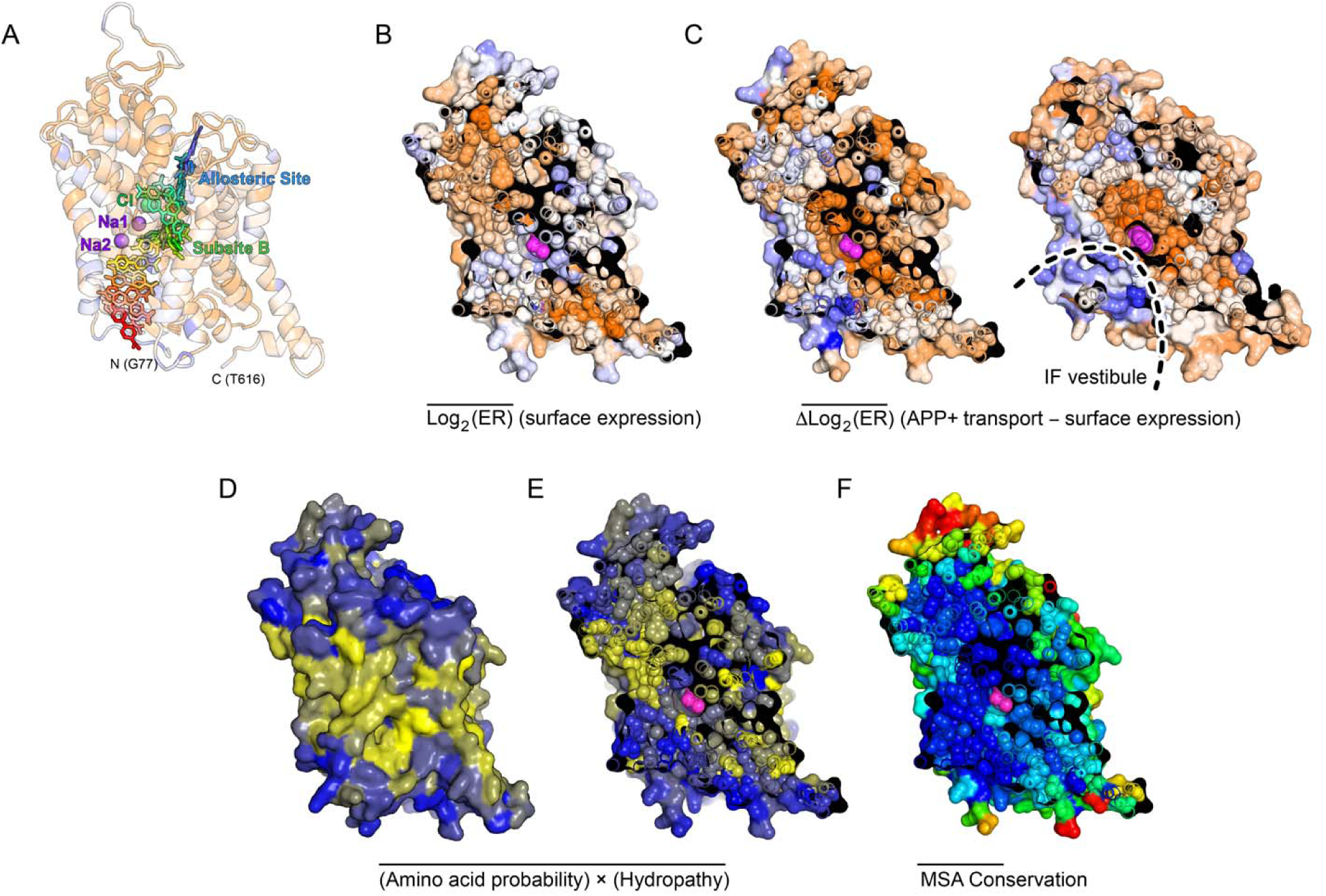
SERT mutations that increase APP+ import are enriched in the intracellular exit pathway. **(A)** Overlaid MD snapshots of APP+ translocation, from when APP+ enters the extracellular vestibule (blue) to its cytosolic exit (red). The positions of ions in the OC state are shown with spheres. SERT is shown as a semi-transparent ribbon and colored by experimental sequence conservation for APP+ transport, showing regions where mutations are depleted in orange to regions where mutations are enriched in dark blue. Also see structure file in Supporting Information online. **(B)** A cross-section through SERT in a partial OF state (PDB 5I73; bound to (*S*)-citalopram in magenta) colored by conservation for folding and surface expression. Tightly conserved residues in the core are orange. **(C)** Cross-sections through SERT in a partial OF state at left and in an ibogaine-bound IF-like state at right (PDB 6DZZ), highlighting in orange residues within the substrate-binding sites and extracellular cavity that become conserved for APP+ transport, while residues that are hot spots for gain-of-transport mutations are dark blue and map to the intracellular exit pathway. Drugs are magenta. **(D, E)** Enrichment ratios for APP+ transport were converted to amino acid probabilities, weighted by hydropathy (yellow, hydrophobic; blue, polar), and averages are mapped onto the **(D)** surface and **(E)** core of SERT (PDB 5I73). Residues lining the entrance cavity and putative exit pathway tolerate polar substitutions. **(F)** Natural conservation based on a sequence alignment of 150 SERT homologs (blue, conserved; to red, variable). Core, entrance, and exit pathways are conserved in natural history.

In the deep mutational scan for APP+ import, the SERT sequence is more conserved than was observed for surface expression alone, with sequence conservation expanded from core regions important for folding towards residues lining the entrance vestibule, the neurotransmitter and ion binding sites, and sites of structural rearrangement near the orthosteric binding site (Figure 2C and 3). For example, a hydrogen bond interaction between Tyr176 and Asp98, which lies ‘above’ the orthosteric binding site, is proposed to serve as a key molecular switch for conformational transitions (*59*). Most mutations to either residue are deleterious for substrate import in the deep mutational scan, which concurs with previous mutational studies of Tyr176 in SERT (*60*), as well as equivalent mutations in DAT and LeuT that exhibit decreased transport activity of their respective substrates (*59*, *61*). Furthermore, the hydrogen bonding network between Glu136, Gly340, and Glu508 (located at the central break in TM6 and forming part of the orthosteric binding site) is critical for mediating the coupling of outward and inward facing states (*62–65*). Previous computational modeling has suggested that the buried Glu136 and Glu508 coordinate a single proton (*66*), while Glu136 also interacts with the Gly340 backbone of TM6. Nearly all mutations to these residues are deleterious for transport function in the deep mutational scan.

**Figure 3.**
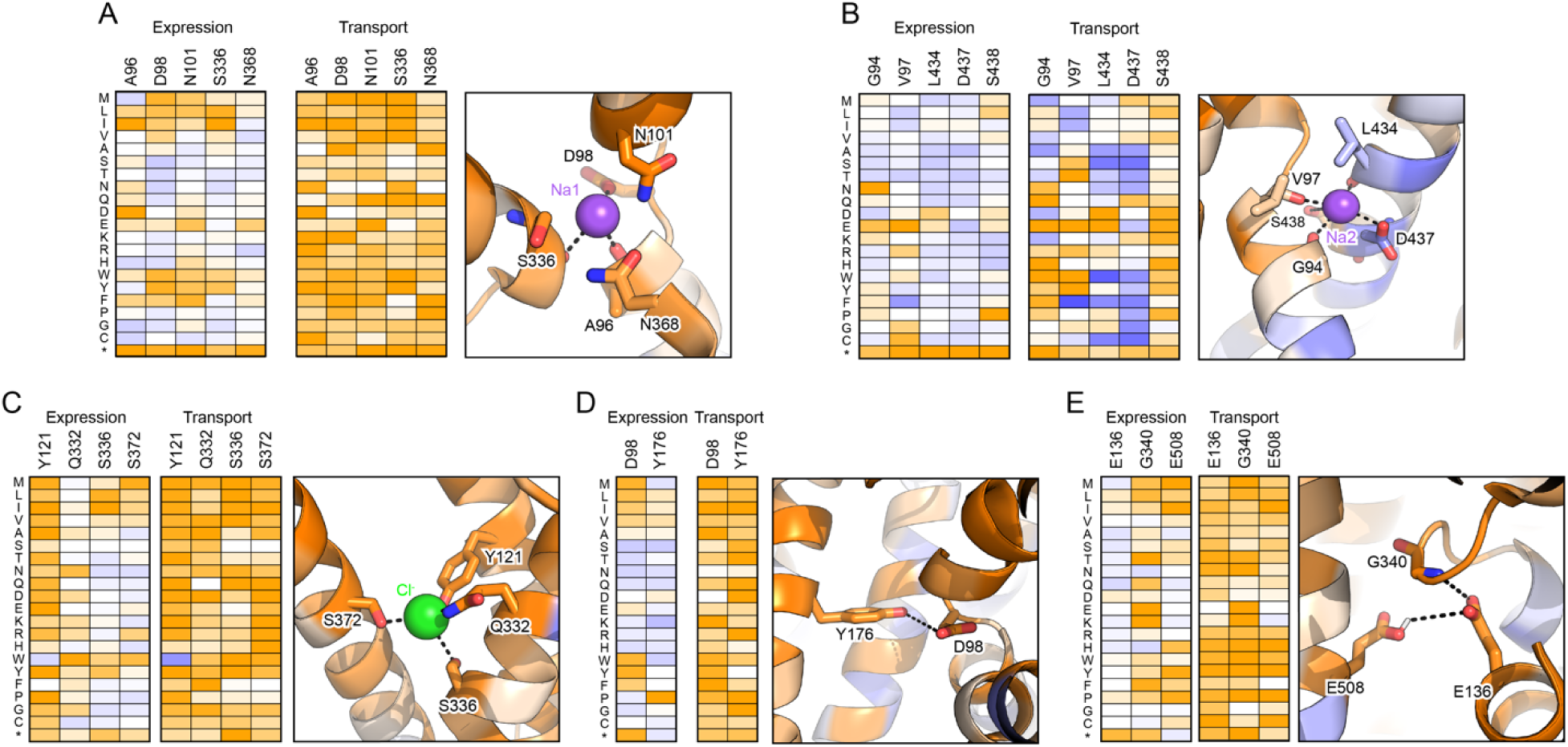
The Na1 site, Cl^-^ binding site, and sites of structural rearrangement adjacent to the orthosteric binding site are conserved for transport function. Heat maps (colored as in Figure 1) show depleted mutations for SERT surface expression (left) or APP+ transport (right) in orange and enriched mutations in dark blue. Accompanying snapshots show SERT (PDB: 5I73) as a cartoon representation with residues colored according to conservation score for transport (as described in Figure 2A). (A) The Na1 site. (B) The Na2 site. (C) The Cl^-^ ion site (modeled based on structural alignment with PDB 7LIA). (D) The hydrogen bonding interaction between Tyr176 and Asp98. (E) The hydrogen bonding network between Glu136, Gly340, and Glu508. Based on previous computational modeling, Glu136 and Glu508 are predicted to coordinate a proton between the two residues (*66*). For this study, Glu136 was modeled in the protonated form in MD simulations.

The mutational scan identifies hundreds of gain-of-function mutations based on positive enrichment after sorting for APP+ import. We further tested by targeted mutagenesis 22 predicted gain-of-function mutations that are spatially dispersed across the structure and validated 21 as trending toward increased APP+ uptake, with 11 having statistical significance (Figure S6). Gain-of-function phenotypes persist when the mutations are introduced into the untagged native cDNA (Figure S2D). We also tested previously reported allelic variants G56A, I425V, and K605N, which have increased 5-HT transport due to altered PKG signaling (*6*, *7*, *67*, *68*), and found these did not have highly elevated APP+ uptake when expressed individually, consistent with their lack of positive enrichment in the mutational scan (Figures S2D and S6A). Cyclic GMP-dependent signaling is therefore unlikely to be an important pathway for modulating SERT activity in the experimental system. Expression of the mutants was generally similar to or less than wild type and cannot explain the large increases in transport activity (Figures S6B and S6C).

Targeted mutagenesis therefore validates predictions from the mutational scan that many amino acid substitutions increase APP+ transport kinetics. Futher validation of the strength of this approach is found in agreement between identified mutants and previous focused studies characterizing the same residues. For instance, our previous study investigating the molecular determinants necessary for transport of 3,4 methylendioxymethamphetamine (MDMA) and 5-HT by SERT (*69*) show that outer gate residue Arg104 is intolerant of substitutions (as also observed in our mutational scan for APP+ import, in which only substitution R104K had a log_2_ enrichment value > −1) and that only acidic residues at Glu493 are permissive for MDMA transport but other substitutions of Glu493 are tolerated for 5-HT (*69*) and APP^+^ transport based on our findings here. Furthermore, residues Ile179 and Asp400 are highly sensitive to substitution in regards to APP^+^ transport, V489 can only tolerate substitution by hydrophobics Leu, Met, Ile, and Phe, and K490 can accept multiple substitutions, which is consistent with the findings of Anderson et al. (*60*).

### Conformational transition to the inward-facing state is rate-limiting for APP+ import

Strikingly, the bulk of mutations that enhance APP+ uptake are clustered in the intracellular half of the transporter (Figure 2C). This clustering of gain-of-function mutations is most noticeable at TM1a, TM6b, and the intracellular segment of TM8, loosely focused near the axis of pseudosymmetry and tightly aligned to the simulated neurotransmitter exit pathway (*70*). After weighting the enrichment ratios by hydropathy and averaging, we find that this region of SERT is highly tolerant of diverse mutations including polar substitutions (Figures 2D and 2E). Substituted cysteine accessability analysis of TMs 1 and 6 by us and others have found 5-HT transport is largely unaffected by cysteine substitutions in intracellular sections of the TMs and accessibility of some of the cysteine substitutions to modifying agents, consistent with their exposure in the transport pathway (*71*, *72*). We propose mutations tolerated in this region will disrupt buried interactions along the exit pathway and thereby facilitate opening of a solvent-exposed intracellular vestibule. That these mutations enhance APP+ uptake is exactly what one would expect if formation of the IF state is rate-limiting.

As previously reported (*44*), the entire Na^+^-coupled import process of the physiological substrate 5-HT has been simulated using Markov state model (MSM) based adaptive sampling (*73–75*). MD with adaptive sampling has been successfully used to simulate complex transport processes in other systems (*74*, *76*). MSM-weighted SERT simulation data are projected onto a coordinate system defined by distances between extracellular and intracellular gating residues (*38*, *44*), thereby visually displaying the relative free energies and barriers between kinetically relevant intermediate states (Figure 4). SERT free of neurotransmitter substrate (i.e. Na^+^-SERT) readily transitions between OF and OC states (Figure 4A) and only in the presence of the cognate substrate 5-HT does the IF state become substantially occupied (Figure 4B), with a reduction in the free energy barrier for the OC-IF transition. Here, the simulations are replicated, but with 5-HT replaced by APP+ in the simulation box as the substrate. Na^+^-bound SERT in an OF conformation obtained from Na^+^-SERT simulations (*44*) was used to seed MD simulations of APP+ import totaling ∼370 μs (referred to as APP-SERT simulations; Figures 4C and S4).

**Figure 4.**
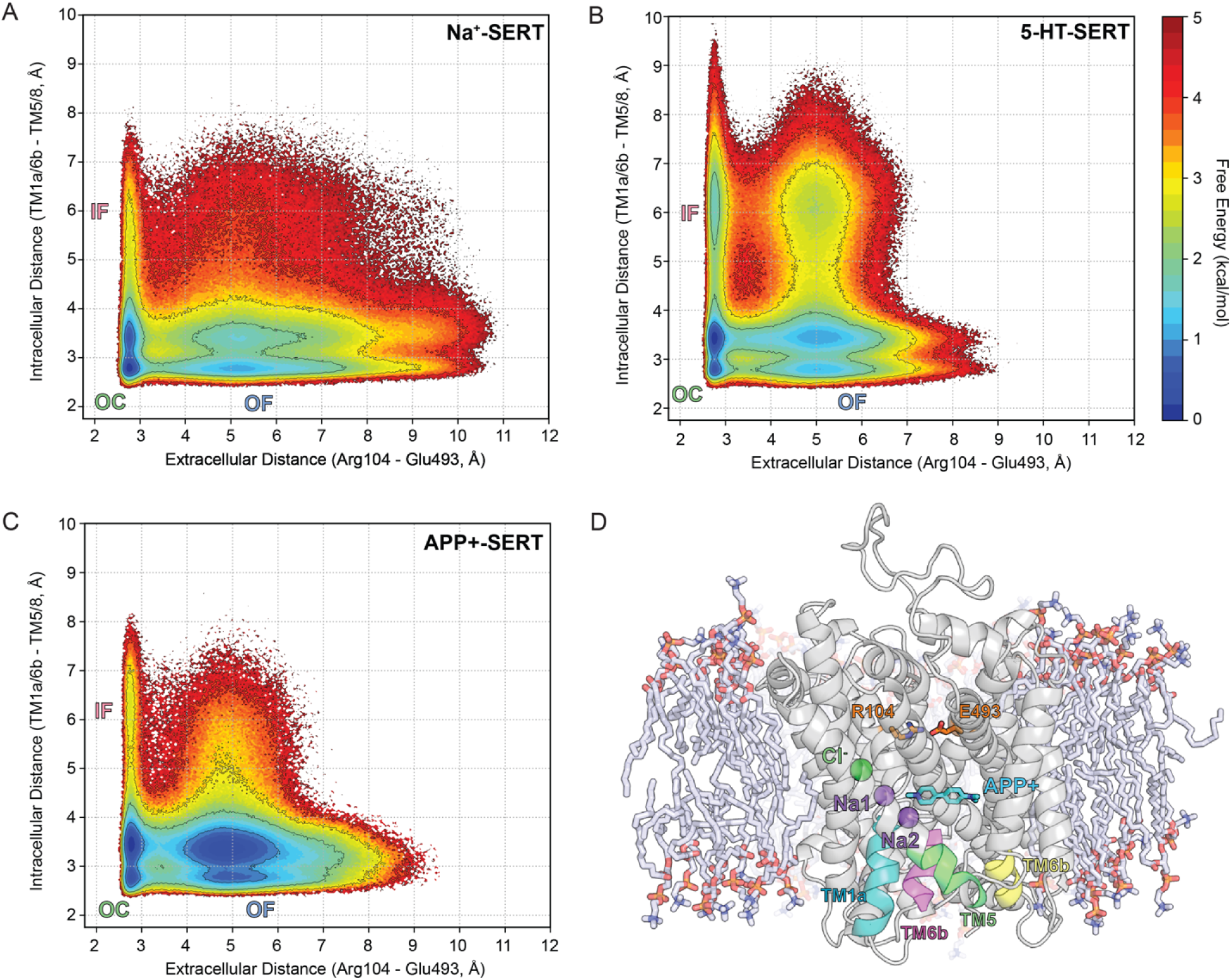
Simulated conformational free energy landscapes of SERT. **(A, B)** Relative free energies from MSM-weighted simulation data plotted against the distances between extracellular and intracellular gating residues for (A) Na^+^-SERT and (B) 5-HT-SERT. These simulation data were previously reported (*44*) and are reproduced here for comparison. The outward-facing (OF) crystal structure (PDB 5I73) was used as the starting structure for MD simulation and transitioned to occluded (OC) and inward-facing (IF) states. **(C)** Calculated free energy landscape of SERT during the APP+ translocation process. **(D)** MD snapshot of membrane-embedded SERT with APP+ bound in subsite B and ions bound in respective sites. Extracellular gating residues Arg104 and Glu493 shown in orange sticks. Intracellular distance was measured based on distances between TM1a, TM6b and the cytoplasmic base of TM5 and TM8, colored in teal, magenta, green, and yellow, respectively.

The simulations reveal that the free energy basin corresponding to IF conformations remains at a higher free energy value in the presence of APP+ (Figure 3C), resembling the conformational free energy landscape for Na^+^-SERT (Figure 3A). The free energy barriers for OC-IF transitions in Na^+^-SERT and APP-SERT are calculated to be 3.43 ± 0.06 kcal/mol and 2.89 ± 0.08 kcal/mol, respectively, which are higher than the estimated 1.66 ± 0.04 kcal/mol barrier for the OC to IF transition when 5-HT is bound and permeating along the transport pathway. Indeed, formation of the IF state is predicted to be rate-limiting for APP+ import, whereas 5-HT interactions facilitate dynamic motions necessary for transport by reducing free energy barriers and stabilizing the IF state (compare Figures 4B and 4C). Similar conclusions regarding differences between substrates have been drawn from experimental observations of bacterial LeuT; only a suitable LeuT substrate such as alanine readily facilitates transition from the OC to IF state (*38*, *77*). The conformational free energy landscape of APP+ import also shows OF states to be further stabilized as the bulkier substrate occupies the extracellular vestibule and the allosteric site. Partial OF-IF-like conformations are also observed, which we previously referred to as an hourglass (HG) state where both gates are open but there is a central constriction (*44*). We speculate that HG conformations may be associated with transient water-conducting states observed in simulations of other membrane transporters (*78*, *79*).

Natural evolution can be considered akin to a deep mutagenesis experiment for protein function as it relates to organism fitness, and we note that residues of SLC6 family members along the permeation pathway are conserved between species (Figure 2F). Furthermore, an exome database shows little human variation (*80*). SERT has therefore reached an optimum (or close to optimum) sequence for its physiological function of ion-coupled 5-HT transport within the constraints of natural evolution. In contrast, our deep mutational scan identifies mutations that increase import of the unnatural substrate APP+ for which the protein sequence is suboptimal. In essence, mutations are found that impose on the APP-SERT conformational free energy landscape the necessary changes that would ordinarily be facilitated by interactions with the native substrate 5-HT.

Based on crystal structures and computational modeling, the N-terminal cytosolic tail of LeuT, in particular LeuT residue Arg5 (equivalent to Arg79 of SERT), forms an interaction network that stabilizes a closed conformation of the cytoplasmic exit cavity in the OF and OC states (*25*, *38*, *41*). The simulations show dynamic motions of the equivalent residues in SERT (a.a. 78-82), with Arg79 forming transient salt bridges to Glu444 and Asp452 to restrict opening of the cytoplasmic exit. Consistent with transitions to the IF state being rate-limiting, mutations to Arg79 and surrounding cytoplasmic gating residues increase APP+ import by destabilizing closure of the intracellular vestibule. We note that alanine substitution of Arg79 (*81*) is deleterious for SERT-catalyzed transport of 5-HT and likely imbalances the conformational equilibrium. Such mutations introduced in the gating residues of GABA transporter GAT-1 have also been shown to alter the equilibria of states and promote inward-facing conformations (*82*). Likewise, mutations in the Na2 site, which are discussed further below, also increase APP+ import but reduce 5-HT transport (*52*). It is only for APP+, which is unable to facilitate appropriate conformational dynamics, that these mutations impart a gain-of-function phenotype, providing a unique mutational data set that functionally maps the substrate exit pathway.

### Deep mutagenesis supports the simulated binding mechanism of Na^+^ and neurotransmitter

The highest flux pathway for ion/APP+ binding events and conformational changes was determined by transition path theory (*83*) and predicts an ordered process for APP+ import identical to that previously simulated for 5-HT (Figure S7). Na^+^ first binds an *apo*-SERT at the Na1 site, followed by a second Na^+^ occupying the Na2 site. APP+ then enters the transporter through the extracellular vestibule and moves to the orthosteric site, with localized dynamics around extracellular loop 4 (EL4) and the extracellular gate, consistent with static structures (*18*, *19*), HDX data (*42*, *43*), and voltage-clamp fluorometry (*84*). Unlike the endogenous substrate 5-HT, the increased structural rigidity of APP+ (Figure S1) results in its translocation ‘down’ the extracellular vestibule to be more constrained, with decreased rotational freedom. In the mutational landscape, hydrophobic substitutions are tolerated or weakly enriched at positions Phe335 and Glu494, where they pack against the APP+ aromatic ring moieties (Figure 5A).

**Figure 5.**
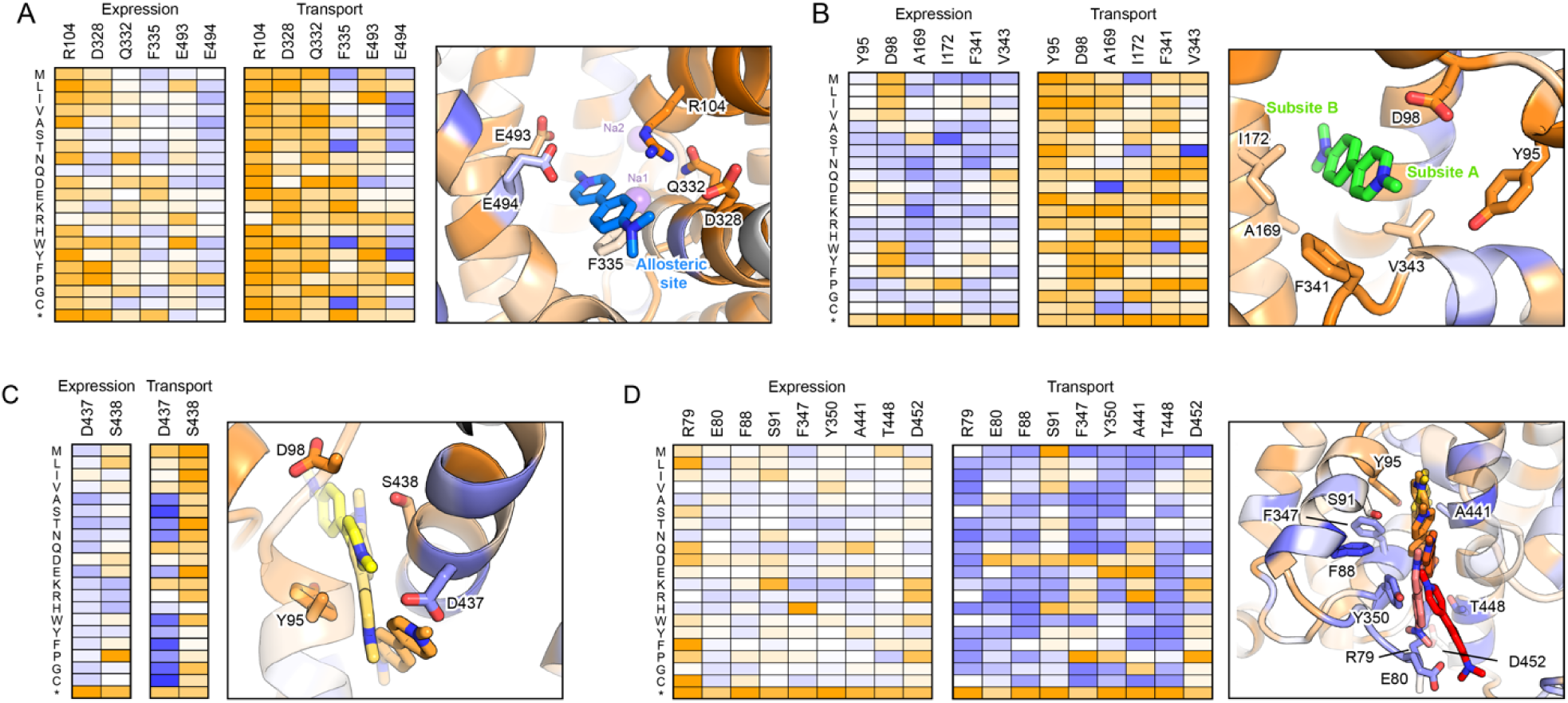
The simulated mechanism for SERT-catalyzed substrate import is supported by the mutational landscape. Heat maps (colored as in Figure 1) show depleted mutations for SERT surface expression (left) or APP+ transport (right) in orange and enriched mutations in dark blue. Accompanying MD snapshots show SERT as a cartoon representation with residues colored according to conservation score for transport (as described in Figure 2A). (A) APP+ entering the transporter via the allosteric site. (B) The neurotransmitter substrate bound to the orthosteric site, straddling subsites A and B. (C) APP+ exiting the orthosteric site through the Na2 site. (D) APP+ permeation along the exit pathway.

Cl^-^, accompanied by an extracellular Na^+^ ion, then engages the extracellular gating residues Arg104 and Glu493, disrupting their electrostatic interactions to allow Cl^-^ to enter the orthosteric cavity. The additional Na^+^ ion does not further progress towards the orthosteric cavity and diffuses back to the extracellular space. Once the neurotransmitter and ions are all bound to their respective sites, the extracellular gates close to form OC conformations. The stability of the OC state promotes the solvation of the intracellular vestibule, thereby weakening the hydrogen bonding network involving various intracellular gating residues (Arg79-Glu452, Glu80-Lys275, Asp87-Trp282, Tyr350-Glu444). Opening of the intracellular permeation pathway is associated with displacement of Na^+^ from the Na2 site into the cytoplasm (consistent with extensive experimental evidence (*36*, *40*, *85*)), followed by cytosolic exit of APP+.

Simulations show contacts made by the neurotransmitter within specific subsites (*32*) inside the orthosteric binding pocket change as import progresses. The aromatic ring of 5-HT shifts from subsite C to B, while flexibility in the aliphatic chain accommodates the amine moiety remaining bound to subsite A (*44*). APP+ never fully straddles subsites A to C, most likely due to steric limitations from having a rigid structure, yet it does interact with subsite B in a binding mode supported by crystallography of dopamine-bound DAT (*30*) and biochemical analyses of inhibitor binding (*32*, *86*, *87*). While residues surrounding the subsites are generally conserved for APP+ import, A169D within subsite B enhances transport (Figure 5B). This mutation is known to also increase the potency of inhibitors (*86*). A169D introduces a carboxylate ∼12 Å distal to where the substrate’s amine group is positioned at subsite A. This spacing is ideal for accommodating the length of APP+, which through resonance has positive charge distributed on both ends for favorable electrostatics in the A169D mutant; this provides further evidence that monoamine substrates are indeed accommodated in subsite B during the transport process. In contrast, Asp in the 169 position has been shown to decrease paroxetine potency via the electrostatic repulsion of the fluorophenyl moiety (*88*).

In the deep mutational scan, mutations to residues coordinating ions in the Cl^-^ and Na1 binding sites are highly deleterious for transport activity (Figure 3A and 3C) and further supports the specific ion dependence for proper function (*52*, *85*, *89*). In contrast, mutations to the Na2 site increase APP+ import, likely due to weakening the interactions between the gating helix TM1 and the scaffold TM8 that would stabilize outward-facing conformations (Figure 3B). Similar mutations introduced to the Na2 site in LeuT biases the conformational equilibria towards the inward-facing state and is demonstrated by its crystal structure (*18*, *85*). In SERT, Na2 site mutations reduce 5-HT binding and transport and underscore the importance of maintaining proper substrate-ion coupling (*52*, *90*).

Gain-of-function mutations that enhance APP+ uptake begin with Leu343 and Asp437 in the Na2 site and continue along the entire exit pathway to the cytoplasm (Figures 3B, 5C, and 5D). There is a near perfect alignment of gain-of-function mutations for APP+ import on TM1a, TM5, TM6b, and TM8 with the simulated permeation trajectory (Figure 4D and structure visualization file in Supporting Information). The Na2 site therefore forms the apex for the cluster of the gain-of-function mutations that delineate the intracellular vestibule. The data are consistent with the simulated release of Na^+^ from the Na2 site, followed by the movement of 5-HT or APP+ into the exit pathway by first passing through the vacated Na2 site. The sharing of a common exit pathway by neurotransmitter and Na^+^ beginning at the Na2 site elegantly explains the 1:1 sodium to neurotransmitter stoichiometry of the transport cycle. Tyr95 forms cation-pi and pi-pi interactions with the substrate and is highly conserved in the mutational scan, tolerating only aromatic substitutions (Figure 5B). The release of 5-HT from the orthosteric site requires a rotameric shift of Tyr95 that is coupled to Na^+^ dissociation at the Na2 site (*44*, *91*). Compared to 5-HT, the structural rigidity of APP+ does not enable Tyr95 to freely rotate and the conformation of Tyr95 is less closely coupled to the release of the substrate (Table S1). Whereas both Na1 and Na2 are important in propagating conformational changes (*66*, *92–95*), the hypothesis that Na^+^ in the Na2 site but not the Na1 site is imported during translocation is further strengthened when considering that members from the SLC5 (eg. galactose transporter) and SLC6 (eg. SERT) families lack sequence homology but share almost superimposable core structures with structurally conserved Na2 sites, whereas only SLC6 members have an Na1 site (*96*) that has been shown to be important for substrate coordination and propagation of conformational changes for translocation (*94*). These distinct roles for Na1 and Na2 are supported by our previous mutagenesis analysis of the Na1 and Na2 sites, where some Na1 site mutants but not Na2 site mutants allowed Ca^2+^ to functionally substitute for Na^+^ in 5-HT transport. Electrophysiological reversal potentials revealed that Ca^2+^ was not transported. These findings along with molecular dynamic simulations supported Ca^2+^ bound to Na1, while not transported, contributes to the coordinated conformational changes for 5-HT transport (*94*).

### Gain-of-function mutations for APP+ uptake at positions outside the permeation pathway are predicted to enhance SERT conformational dynamics

We clustered the SERT mutational data for surface expression and APP+ transport activity using an unsupervised learning algorithm, Uniform Manifold Approximation and Projection (UMAP) (*97*). UMAP presents an approach to effectively reduce the high dimensional mutational landscape and systematically derive quantitative relationships between a residue’s mutational response and its biophysical and biochemical properties (*98*, *99*). Visualization of the mutational landscape on the two-dimensional UMAP space shows that residues cluster based on various biochemical and biophysical properties (Figures 6 and S5).

**Figure 6.**
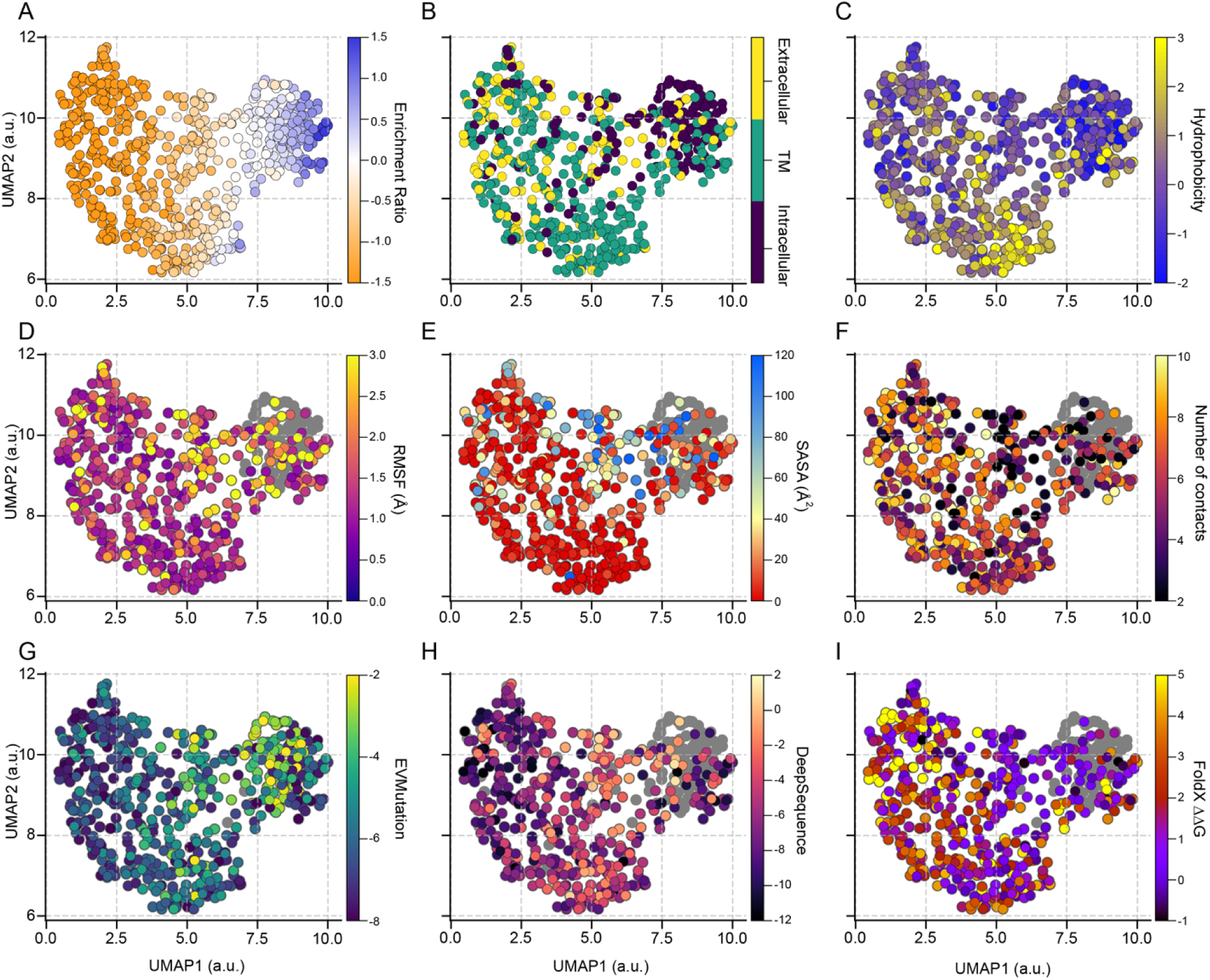
UMAP projection of the mutational landscape for SERT-catalyzed APP+ transport. Each residue was clustered using the UMAP clustering algorithm based on the activities of all amino acid substitutions at that position, and are colored based on different biochemical and biophysical properties: (**A**) APP+ uptake enrichment ratio, (**B**) residue location, (**C**) Kyte-Doolittle hydrophobicity, (**D**) root mean square fluctuation (RMSF) (**E**) solvent accessible surface area (SASA), and (**F**) average number of contacts. The mutational landscape was also compared to scores from variant effect predictors (**G**) EVmutation, (**H**) DeepSequence, and (**I**) FoldX. Residues in which data is unavailable are colored gray. The first UMAP dimension strongly correlates with enrichment ratio (transport ρ = 0.966). The properties from MD simulations (RMSF, SASA, and average number of contacts) were calculated on 50,000 structures drawn from an MSM-weighted distribution. Projection of the SERT expression deep mutational landscape in 2D UMAP space is available in Supplemental Information Figure S8. Correlations between biochemical and biophysical properties with the UMAP dimensions and DMS enrichment ratios are listed in Table S2.

Projection of the mutational landscape for APP+ transport shows that residues naturally cluster based on residue site location (Figure 6B) and hydrophobicity (Figure 6C). When compared to biophysical properties obtained from MD simulations, we observed that a residue’s conservation score in the deep mutational scan is related to its dynamic fluctuations (Figure 6D), solvent accessible surface area (SASA) (Figure 6E), and local structural packing (Figure 6F). Residues that enhance transport activity are associated with larger fluctuations, while residues with low mutational tolerance exhibit low SASA and RMSF, suggesting these residues are buried in the protein core and are important for stabilizing contacts to maintain proper folding. Furthermore, the deep mutational landscape for SERT expression correlates with mutational effects predicted by EVmutation (*100*) (Figure S8G), DeepSequence (*101*) (Figure S8H), and FoldX (*102*) (Figure S8I), in addition to correlating with local structural packing (Figure S8F), underscoring that sequence and structural constraints are necessary for proper expression and trafficking to the plasma membrane (Table S1). By comparison, the mutational landscape for APP+ transport is less correlated with the variant effect predictions, further emphasizing that important protein features are only partially shared between the transport mechanisms for the endogenous and nonendogenous substrates.

While less abundant, some mutations outside the simulated permeation pathway are also enriched for APP+ import, including mutations at helical hinges that flex during structural transitions (e.g. A173G, P499K, S585P), or on neighboring helices immediately adjacent (e.g. F263C, P499K, G582K); these substitutions may stabilize unwound structural features within the membrane phase, facilitate dynamic motion, alter packing against surrounding helices, or stabilize conformational states. The mutational tolerance of Ala173 at a helix kink is particularly striking; nearly all substitutions are deleterious for transport except for the small amino acid glycine that can increase flexibility (Figure 7B). DAT and NET both have glycine at this position. We also find a cluster of gain-of-function mutations for APP+ import within the hydrophobic core where TM1b packs against EL4 (Figure 7C); substantial EL4 motion in bacterial transporters can occlude the extracellular vestibule (*103*), while more limited EL4/TM1b motions are observed in cryo-EM structures of ibogaine-bound SERT (*19*) and in the simulations. In general, mutations in this region that increase APP+ import substitute large hydrophobics for smaller side chains that reduce packing and are therefore anticipated to increase dynamics, possibly creating localized molten globule-like behavior.

**Figure 7.**
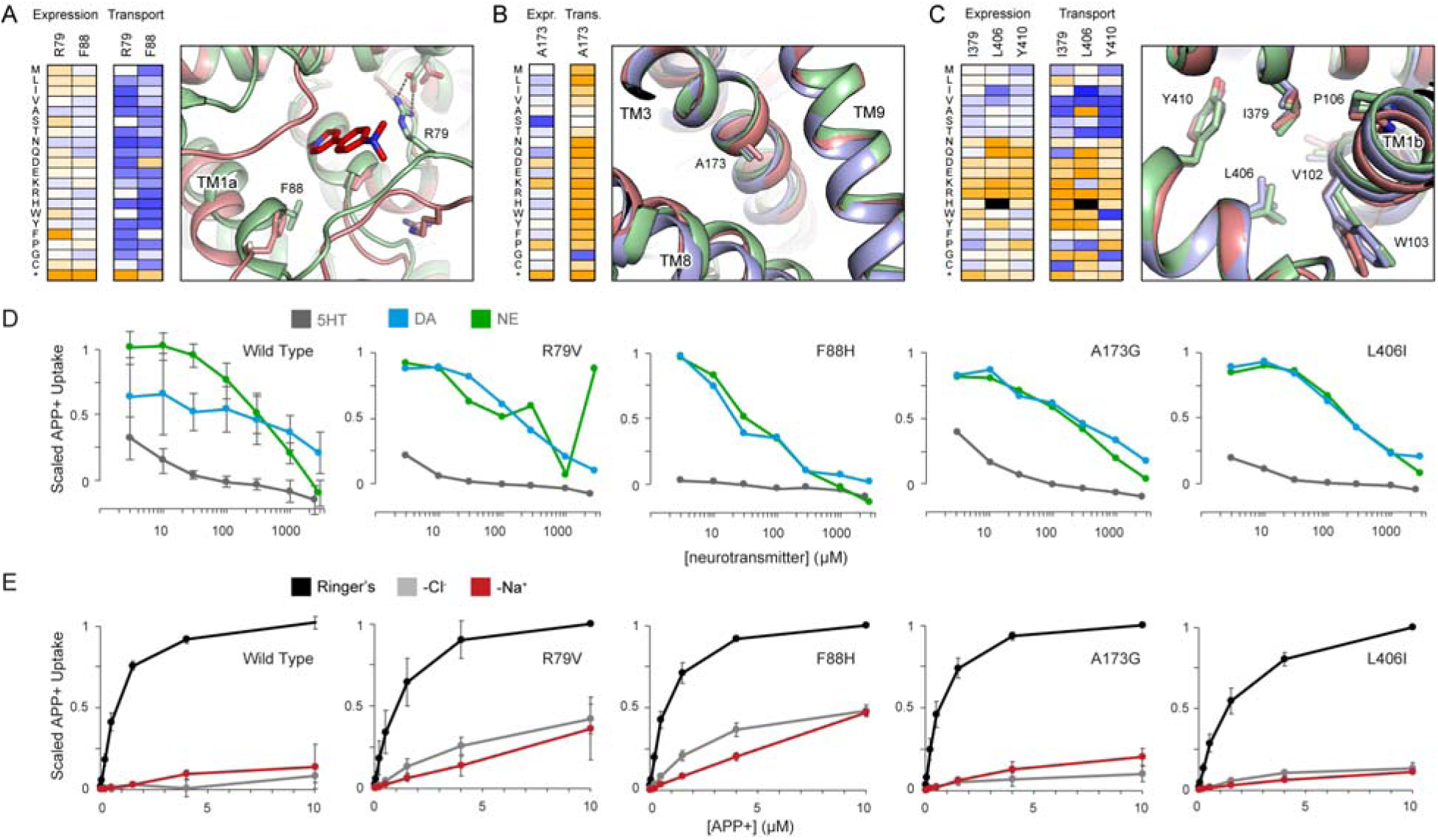
Exit pathway mutations can cause a partial loss of Na^+^ and Cl^-^ dependency. **(A-C)** Superimposed MD snapshots (blue, OF state; green, OC state; and pale red, IF state) showing dynamic regions. Experimental enrichment ratios for select residues are plotted alongside, colored orange (deleterious mutations) to dark blue (enriched mutations) as in Figure 1. **(A)** Motion around the intracellular gate. **(B)** A hinge in TM3. **(C)** The hydrophobic core below EL4, adjacent to the extracellular gate. **(D)** APP+ uptake in SERT-expressing cells was more potently inhibited by 5-HT (grey) than by norepinephrine (NE, green) or dopamine (DA, blue). (Mean; WT n = 5, R79V n =2, F88H n =2, A173G n = 3, L406I n = 2. For WT, error bars show SD.) **(E)** APP+ uptake by SERT-expressing cells was measured in Ringer’s solution (black), with replacement of Na^+^ (red) or Cl^-^ (grey) ions. Transport was Na^+^ and Cl^-^ dependent for WT, A173G, and L406I SERT, while showing a partial loss of ion dependence in R79V and F88H mutants. (WT n = 4, R79V n = 4, F88H n = 3, A173G n = 4, L406I n = 3, mean ± SD)

We further investigated four SERT mutations that increase APP+ import, chosen to represent different structural regions: R79V in the cytoplasmic gating loop (Figure 7A), F88H on TM1a proximal to the exit pathway (Figure 7A), A173G at a helix kink in TM3 (Figure 6B), and L406I in EL4b where it packs against TM1b (Figure 7C). All mutants retained selectivity for 5-HT over dopamine and norepinephrine (Figure 7D). Na^+^ and Cl^-^ dependency of APP+ import is partially lost in two of the mutants, R79V and F88H (Figure 7E), which both lie on the exit pathway and are predicted to destabilize closure of the intracellular vestibule. Correct ion coupling for symport therefore demands appropriate free energy differences between conformational states, ensuring that all ions and substrate must bind to energize structural transitions.

To further support the conclusion that SERT gain-of-function mutations for APP+ import in the deep mutational scan are substrate specific, we evaluated the set of four SERT mutants for transport of tritiated 5-HT in transfected HEK-293 cells (Table 1). SERT A173G and L406I have similar kinetic parameters for 5-HT transport as the wild type, albeit with slightly reduced maximum velocity (V_max_). However, SERT mutants R79V and F88H have large decreases in V_max_ and K_M_, possibly due to loss of interactions with 5-HT in the intracellular exit pathway that would ordinarily facilitate translocation out of the orthosteric site and down a narrow intracellular vestibule toward bulk solvent (*44*). Others have also found that mutations that favor IF conformations lower both V_max_ and K_M_ (*57*).

## DISCUSSION

Protein sequences are highly optimized over natural history for their physiological functions. Here, we employed a fluorescent-based assay to determine the effects of nearly all amino acid substitutions on human SERT surface expression and transport. The selection of SERT mutants for surface expression reveals conservation and evolutionary constraints for plasma membrane trafficking and demonstrates the necessity for maintaining the proper three-dimensional fold, rather than just membrane helix insertion and biogenesis per se. Furthermore, SERT variants were selected for enhanced transport of an unnatural substrate for which the protein sequence is suboptimal, and which simulations indicate is unable to promote the conformational changes that drive transport. As a consequence, hundreds of gain-of-function mutations were discovered that richly inform molecular mechanism and provide insight into conformational changes that limit transport kinetics. We conclude that in the absence of the cognate substrate, there is a high free energy barrier for transition to the IF state, which will maintain ion-neurotransmitter coupling and prevent ion leakage, and the symported Na^+^ and neurotransmitter share a common exit pathway that begins with the Na2 site at the apex of the intracellular vestibule. This proposed import mechanism is likely shared with related neurotransmitter:sodium symporters.

The technical requirements for *in vitro* selection restricted investigations to the fluorescent substrate and neurotransmitter analogue APP+ (*104*), which is larger than and has reduced configurational entropy relative to 5-HT (Figure S1), and its import was therefore sensitive to mutations along the permeation pathway. The concept of discovering mechanism by studying an alternative substrate is widely practiced in the field of enzyme directed evolution (*105*), and engineering of major facilitator superfamily transporters to redirect substrate preference has illuminated important motifs and sequence features (*106–109*). Notably amongst studies of NSS homologues, rational mutagenesis switched LeuT from being inhibited by to now transporting tryptophan, demonstrating that substrate complementarity is necessary for driving the OF-OC structural transition (*110*). Our work here similarly uses an unnatural substrate to inform the mechanism of monoamine NSS transporters through the discovery of gain-of-function mutations.

SERT self-associates into poorly defined oligomers containing up to 8 subunits (*111*, *112*), consistent with aggregation or lipid microdomain localization as opposed to distinct protein-protein interactions. The structures of human SERT and *Drosophila* DAT are both monomeric (*19*, *26*, *30*), and while detergent extraction can dissociate protein-protein interfaces, it nonetheless suggests against a specific, tight oligomer. Crystal structures of LeuT obtained in a dimer suggested that the two monomers may form an interface at TM12 (*18*). Periole et al. conducted coarse-grained SERT simulations and observed the probability of various dimer interfaces involving different transmembrane helices, of which symmetric TM12-TM12 and asymmetric TM7-TM12 dimers were most favorable (*113*). In light of these studies, it remains unknown whether SERT forms specific oligomers that are biologically relevant. In investigations of a dimeric G protein-coupled receptor that is closely related to metabotropic neurotransmitter receptors, deep mutagenesis identified membrane-exposed surface patches that were relatively highly conserved during *in vitro* selection; the data guided the integrative structure determination for the protein’s dimeric architecture that closely agreed with cryo-electron microscopy (*46*). Searching for surface patches on SERT that are conserved in the deep mutational scans, the landscapes for SERT expression and APP+ transport identify the membrane-exposed surface formed by TM2 and TM7 as a candidate oligomerization or lipid-interaction site. However, these helices also form critical contacts to Cl^-^ and Na^+^ within the protein core, and their higher conservation may simply reflect their importance for proper folding with bound ions. The mechanisms by which SERT proteins self-associate requires further investigation.

The mechanistic details described here may provide insights into how SERT kinetics are regulated *in vivo*. For example, syntaxin 1A interacts with the SERT N-terminus to dampen excess Na^+^ flux during 5-HT transport (*23*, *114*), and it may be that syntaxin 1A stabilizes the OF state with a bound N-terminal gating loop to ensure free energy barriers are sufficiently high for strict coupled transport. By comparison, phosphorylation of Thr276 by PKG when SERT is in the IF conformation increases transport velocity, possibly by preferentially stabilizing the IF state (*68*). Hence under certain physiological conditions, nature may have exploited features of the conformational free energy landscape to increase kinetics or change ion coupling and flux.

## METHODS

### Cell Culture

Expi293F cells (ThermoFisher) were cultured in Expi293F expression medium (ThermoFisher) at 37°C, 8% CO_2_, 125 rpm. GripTite HEK-293 (ThermoFisher) cells were grown in Dulbecco’s Modified Eagle Medium supplemented with 10% fetal bovine serum and 150 ng/ml G418.

### Reagents

APP+ was from Aobious, prepared as a 100 mM solution in DMSO, and stored at −80°C. Neurotransmitters 5-HT, norepinephrine, and dopamine were obtained from Sigma-Aldrich and prepared as 100 mM (dopamine and norepinephrine) or 25 mM (5-HT) solutions in water. Neurotransmitter stocks were stored at −80°C until use.

### Plasmid Construction

Codon optimized human SERT was assembled from synthetic DNA fragments (IDT) and inserted into the NheI-XhoI sites of pCEP4 (Invitrogen). The use of a synthetic gene ensures there are no sequence features, such as GC-rich tracts, resistant to PCR-based mutagenesis, and therefore aids in library construction. A c-myc epitope tag flanked by a glycine / serine rich linker was inserted into EL2 between residues 216 and 217. A version without the tag was created by PCR-based mutagenesis. For protein expression from the native human cDNA, SERT was subcloned from pcDNA3-hSERT (Addgene 15483)^68^ into Kpn1-Xho1 sites of pCEP4. A c-myc epitope tag was inserted into EL2 using overlap extension PCR. To measure surface expression of SERT normalized by total expression, superfolder GFP was fused directly to the myc-SERT N-terminus. All constructs featured a consensus Kozak sequence. Important plasmids are deposited with Addgene.

### Library Construction

Two separate SSM libraries were created from plasmid pCEP4 encoding codon optimized myc-SERT. The SERT N-terminal library covered amino acids 2-325, and the C-terminal library covered amino acids 326-630. For each library, overlap extension PCR using primers containing degenerate NNK codons was applied (*115*), and the pooled PCR products were cut and ligated into NheI-NotI or HindIII-XhoI sites for the N-terminal and C-terminal libraries, respectively. Ligation products were electroporated into NEB5α cells (New England Biolabs), with the number of transformants at least 50 times the possible DNA sequence diversity. Plasmid libraries were purified using GeneJet Maxi Prep kit (ThermoFisher).

### Library Selection

Libraries were expressed transiently using Expifectamine (ThermoFisher) in Expi293F cells. To reduce the likelihood of two SERT sequences being acquired by a single cell (*48*), 1ng library DNA was diluted with 1.5 μg pCEP4ΔCMV (*46*) carrier plasmid per milliliter of culture at 2×10^6^ cells/ml. Cells were incubated at 37 °C, 8% CO_2_, with continuous shaking at 125 rpm for 2 h, followed by media replacement, and then incubated for a further 22 h. Transport activity was assessed by washing with PBS-BSA (phosphate-buffered saline supplemented with 0.2% bovine serum albumin), and incubating in 0.5 μM APP+ (Aobious) in PBS-BSA for 15 min at room temperature with gentle agitation. Cells were then washed with ice cold PBS-BSA, incubated on ice for 20 m with 1/200 anti-myc-Alexa 647 (clone 9B11; Cell Signaling Technology), washed twice, and then analyzed on an Accuri C6 or BD LSRII cytometer, or sorted on a BD FACS ARIA II. During sorting, the main cell population was gated by forward scatter-A versus -W to exclude doublets and debris. Propidium iodide-negative events were then gated to remove dead cells. The top 50 % of SERT-expressing cells (based on Alexa 647 fluorescence) were collected and frozen at −80 °C. For analysis of transport, the top 10-15 % of APP+ fluorescent cells were collected within the SERT-expression gate. To adequately sample each mutation in the libraries during sorts, at least 100,000 cells were collected, pooling frozen cell pellets from separate days as necessary to create one sample from which cDNA was prepared. Typically, ∼8 h of sorting was required for each condition, and transfected cultures were not sorted for longer than 4 hours to maintain viability. Each deep mutational scan experiment consisted of at least two separately sorted and prepared transfected cell samples.

### Library Sequencing and Analysis

Total RNA was extracted from the sorted cells (GeneJet kit from ThermoFisher), followed by first strand cDNA synthesis primed with gene-specific oligonucleotides using high-fidelity Accuscript (Agilent). DNA fragments were amplified in two rounds of PCR as previously described (*46*, *115*), adding Illumina adapter sequences and barcodes. Samples were sequenced on an Illumina MiSeq or HiSeq2500 using 2×250 nt paired end read protocol. Data were processed using Enrich (*116*), and commands for analysis are included in the GEO submission. Enrichment ratios for each variant were calculated by dividing the transcript frequency from the selected population by the plasmid DNA frequency in the naive plasmid library. Sequence variants with < 10 reads (equivalent to a frequency of ∼3.5 x 10^-6^) in the naïve plasmid library were considered missing. Log_2_ enrichment scores are displayed as heat maps using gnuplot (http://www.gnuplot.info/), or displayed on the crystal structure of (s)-citalopram-bound SERT (PDB 5I73) using PyMOL (Shrodinger, LLC).

### Variant Selection, Targeted Mutagenesis, and Analysis

Variants for validation by targeted mutagenesis were selected from the deep mutational scan with a log_2_ enrichment ratio of >3 for APP+ import and <1 for expression, or were reported human alleles. Primers containing the specific codon change were used for site-directed mutagenesis using overlap extension PCR. SERT variants (500 ng plasmid DNA per ml culture) were transiently transfected into Expi293F cells (2×10^6^ cells/ml) using Expifectamine (ThermoFisher). Transfection enhancers were added 16-18 h post-transfection, and cells were harvested for analysis 36 h post-transfection. APP+ transport activity was assessed as described for library preparation. Samples were analyzed on an Accuri C6 cytometer. Total and surface protein expression were measured using the BD Fixation/Permeabilization Solution Kit (BD Biosciences) in conjunction with anti-myc-Alexa 647 staining (clone 9B11; Cell Signaling Technology). For testing neurotransmitter selectivity, assays were performed on Expi293F cells incubated with 0.5 μM APP+ and varying concentrations of 5-HT, dopamine, or norepinephrine in Ringer’s Solution (140 mM NaCl, 5 mM KCl, 1.2 mM MgCl_2_, 5 mM Glucose, 2 mM CaCl_2_, 10 mM HEPES, pH 7.5) supplemented with 0.2% bovine serum albumin. Ion dependency was measured by replacing NaCl in Ringer’s-BSA solution with Choline-Cl (140 mM) and adjusting the pH to 7.5 with choline-hydroxide solution, or by replacing chloride salts with salts of gluconate, and adjusting pH with KOH.

The mean fluorescence (*F_APP+_*) of cells incubated with APP+ was measured by flow cytometry. Background autofluorescence (*F_0_*) was subtracted using SERT-transfected cells without APP+ to calculate *ΔF = F_APP+_ – F_0_*. To assess transport activity (*T*), *ΔF* of mock transfected control cells incubated with APP+ was subtracted from *ΔF* of SERT-expressing cells: *T = ΔF_SERT_ – ΔF_negative_*. For quantifying surface expression, autofluorescence of untransfected cells incubated with Alexa 647-conjugated anti-myc antibody was subtracted from the mean fluorescence units of the SERT-transfected population. All replicates are independent samples and statistical tests as described in legends were calculated using Prism 10.

### [^3^H] 5-HT Single-Point Uptake Assay

GripTite HEK-293 cells were transiently transfected with 100 ng of plasmid DNA and grown in 24-well CulturPlate-24 plates (PerkinElmer) to 80-90% confluence. Cells were washed twice with 500 μL of 37 °C Modified Krebs-Ringer HEPES + glucose (MKRGH: 5 mM Tris, 7.5 mM HEPES, 120 mM NaCl, 5.4 mM KCl, 1.2 mM CaCl_2_, 1.2 mM MgSO_4_, 10 mM glucose, pH 7.4). Uptake was performed in a final volume of 250 μL with 20 nM hydroxytryptamine creatinine sulfate, 5-[1,2-^3^H(N)] ([^3^H]5-HT) at 37.5 Ci/mmole (PerkinElmer, Cat# NET498001MC). Uptake was allowed to proceed for 10 min at 37 °C followed by three washes with ice-cold MKRH, aspirated dry, and solubilized with 250 μL of MicroScint-20 (PerkinElmer). [^3^H]5-HT uptake was measured with TopCountNXT Microplate Scintillation and Luminescence Counter (PerkinElmer) and nonspecific uptake determined with 10 μM (*S*)-citalopram treated and parental cells.

### Saturation Uptake Analysis

GripTite HEK-293 cells were grown, transfected, and washed as above. 200 μl of MKRHG was as added to each well followed by 2-fold serial dilutions of non-radiolabled 5-HT prepared in MKRHG with ascorbic acid and iproniazid. [^3^H]5-HT was added to each well to a final concentration of 30 nM. After 2.5 minutes, the cells were washed three times with ice-cold MKRH and radioactivity quantified (CPM) as above. CPMs were converted to amoles based on specific activity of 5-HT in each well and counter efficiency. The rate of uptake was plotted versus 5-HT concentration to determine K_M_ and V_max_.

### All Atom MD Simulations

Simulations were initiated from an OF Na^+^-bound SERT structure, with Na^+^ bound in the Na1 and Na2 sites, obtained from our previous study (*44*). The protein was embedded in a 1-palmitoyl-2-oleoyl-sn-glycero-3-phosphocholine (POPC) bilayer using the Membrane Builder plugin in CHARMM-GUI (*117*). The terminal chains were capped with acetyl and methyl amide groups. 1 APP+ molecule was added to the simulation box. 150 mM NaCl was added to neutralize the system and mimic physiological conditions. Glu508 was modeled in its protonated form. The disulfide bond between Cys200 and Cys209 of EL2 was present. The system was solvated with TIP3P water molecules (*118*) in a box volume of 75 x 75 x 112 Å^3^. Simulations were implemented with the Amber18 package (*119*) using Amber ff14SB(*120*) and GAFF2 force fields(*121*). Parameters for the APP+ molecule were generated from the Antechamber package (*122*). The MD system was minimized for 20,000 steps using the conjugate gradient method, heated from 0 to 300 K at NVT, and equilibrated for 40 ns under NPT conditions. All production runs were conducted at constant NPT conditions (300K, 1 atm), periodic boundary conditions, and hydrogen mass repartitioning (*123*). An integration timestep of 4 femtosecond (fs) was used. Temperature (300 K) was maintained using Langevian dynamics with a 1.0 picosecond^-1^ damping coefficient. Pressure (1 atm) was maintained using Monte Carlo barostat with an update interval every 100 simulation steps. The nonbonded distance cutoff was set at 10 Å. Electrostatic interactions were treated with the Particle Mesh Ewald method (*124*), and hydrogen bonds were constrained using SHAKE algorithm (*125*). Simulation snapshots were saved for every 100 ps during production simulations.

As conducted in our previous study, we implemented a Markov state model (MSM)-based adaptive sampling approach (*126*, *127*) to thoroughly sample the conformational landscape of SERT during the APP+ import process. This approach involved an iterative sampling procedure in which multiple simulations were conducted in parallel. Data were then clustered based on a designated metric of the z-position of the symported Na^+^, Cl^-^, and APP+ using a K-means algorithm. States from the least populated clusters were chosen for the next iteration of simulations. A total of ∼370 μs of simulation time was collected and thoroughly sampled the conformational landscape (Figure S9A).

### MSM Construction

Markov state modeling is a statistical approach for discretizing the simulation data into kinetically relevant clusters and calculating the transition probability between each cluster (*73*). MSM were constructed with the pyEMMA 2.5.6 Python package (*128*). To compare the APP+ import process with Na^+^ and 5-HT-SERT simulations, we constructed the MSM using the same 16 interhelical distances surrounding the channel pore radius and z-components of substrates (APP+, Cl^-^, and the symported Na^+^) as featurization metrics (Figure S10A). The number of clusters was optimized based on the VAMP scoring function implemented in pyEMMA. 4 time independent components (tICs) and 600 clusters were chosen to construct the MSM (Figure S10B). The lag time of 12 ns was determined to be a Markovian lag time based on implied timescale plots (Figure S10C). To validate the model, we perform a Chapman-Kolmogorov test (*129*) on a 4 macrostates MSM (Figure S11).

### Trajectory Processing and Analysis

The CPPTRAJ module (*130*) implemented in AmberTools and MDTraj (*131*) was used for post processing of trajectory data. All simulation data were projected on the coordinate landscape defined by the extracellular gating distance as the closest heavy atom between Arg104 and Glu493 and the intracellular distance as the closest heavy atom between the groups of residues Val86-Ser91, Tyr350 and Val199-Trp282, Glu444. To estimate the error of the free energy landscape, a bootstrapping method was implemented in which 500 independent MSM were constructed using a random selection of 80% of the trajectory dataset (Figure S9B). Trajectories and MSM states were visualized with Visual Molecular Dynamics (VMD) (*132*) and PyMOL (Shrodinger, LLC). In-house scripts and the matplotlib python package were used to generate plots.

### Clustering the mutational landscape using UMAP

The activities of all amino acid substitutions per SERT position were clustered using UMAP python package (github.com/lmcinnes/umap) using the following hyperparameters: n_neighbors = 10, min_dist = 0.2, n_components = 2, metric = euclidian. The mutational data were then projected on the two UMAP components and colored based on different biochemical and biophysical properties. The hydrophobicity of residues was calculated using the Kyte-Doolittle scale using a window size of 9. Features from MD simulations, root mean square fluctuations (RMSF), solvent accessible surface area (SASA), and average number of residue-residue contacts, were calculated on 50,000 structures drawn from an MSM-weighted distribution. The outward-facing SERT structure (PDB: 5I73) was used as the reference structure for RMSF calculation. SASA calculation was implemented using the MDtraj package and a 1.4 Å probe radius. An 8Å Cα-Cα distance cutoff was used to determine residue-residue contacts. Variant effect predictions were generated by EVmutation (github.com/debbiemarkslab/EVcouplings) and DeepSequence (github.com/debbiemarkslab/DeepSequence) packages and used the EVcouplings package to construct the input sequence alignment. *In silico* free energy changes as a result of a mutation were predicted using the FoldX suite (foldxsuite.crg.eu).

### Estimating Evolutionary Conservation

Natural sequence conservation was mapped to the SERT structure (PDB 5I73) using the ConSurf server (*133*). Homologs were searched using CS-BLAST from UNIREF-90 with an E-value cutoff of 0.0001 and 3 iterations. Maximum and minimum identity were set at 95 % and 35 %, respectively. 500 NSS transporter sequences were aligned with MAFFT and conservation scores calculated using Bayesian algorithm of substitution.

### Data and Software Availability

Raw deep sequencing data and calculated enrichment ratios are deposited with NCBI’s Gene Expression Omnibus under series accession number GSE109499. Command lines for analysis with Enrich are also included. Representative structures from the Markov state model and MD simulations are available on Zenodo at https://doi.org/10.5281/zenodo.4695032.

## SUPPORTING INFORMATION

Online Supporting Information includes:

PDF of Supplementary Figures S1 to S8

PyMOL file of SERT colored by conservation score for APP+ transport with overlaid APP+ permeation pathway

Movie S1 – morph of SERT conformational transitions colored by APP+ transport conservation score

Excel file of enrichment ratios from the SERT deep mutational scans

## AUTHOR INFORMATION

### Corresponding Authors

- (E.P.)
- (D.S.)

### Author Contributions

H.J.E. and S.K.S. did experimental characterization and deep mutagenesis. M.C.C. and B.S. did MD simulations. E.W. and L.K.H. tested uptake of 5-HT. E.P., L.K.H., and D.S. supervised and directed research.

### Notes

The authors declare they have no conflict of interest. E.P. receives stock and compensation from Cyrus Biotechnology for working on drug programs unrelated to this study.

## ACKNOWLEDGEMENTS

Within the UIUC Roy J. Carver Biotechnology Center, Barbara Pilas and Angela Kouris assisted with flow cytometry, and Alvaro Hernandez and Chris Wright assisted with deep sequencing. We thank Jesse R. Horne for technical assistance on DeepSequence and FoldX calculations. This work was funded by R21 MH113155 from NIMH to E.P., and D.S. is supported by NSF Early Career Award by NSF MCB 18-45606. The research is part of the Blue Waters sustained-petascale computing project, which is supported by the National Science Foundation (awards OCI-0725070 and ACI-1238993) and the state of Illinois, and as of December, 2019, the National Geospatial-Intelligence Agency. Blue Waters is a joint effort of the University of Illinois at Urbana-Champaign and its National Center for Supercomputing Applications.

