## Supplementary Figures & Tables for "Deep Mutagenesis of a Transporter for Uptake of a Non-Native Substrate Identifies Conformationally Dynamic Regions"

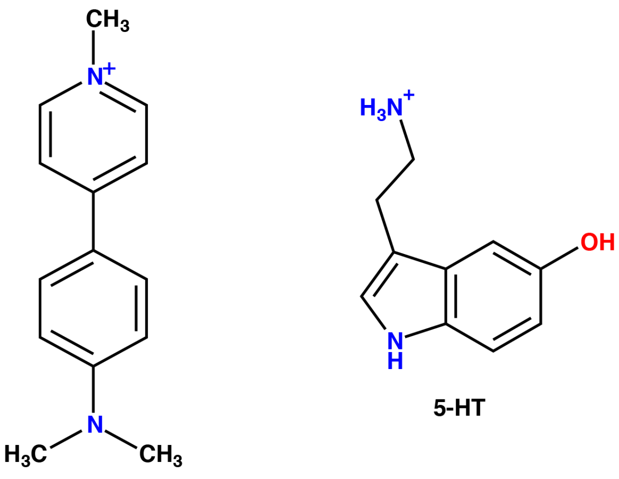

APP+

**Figure S1. Chemical structures of APP+ and 5-HT (serotonin).**

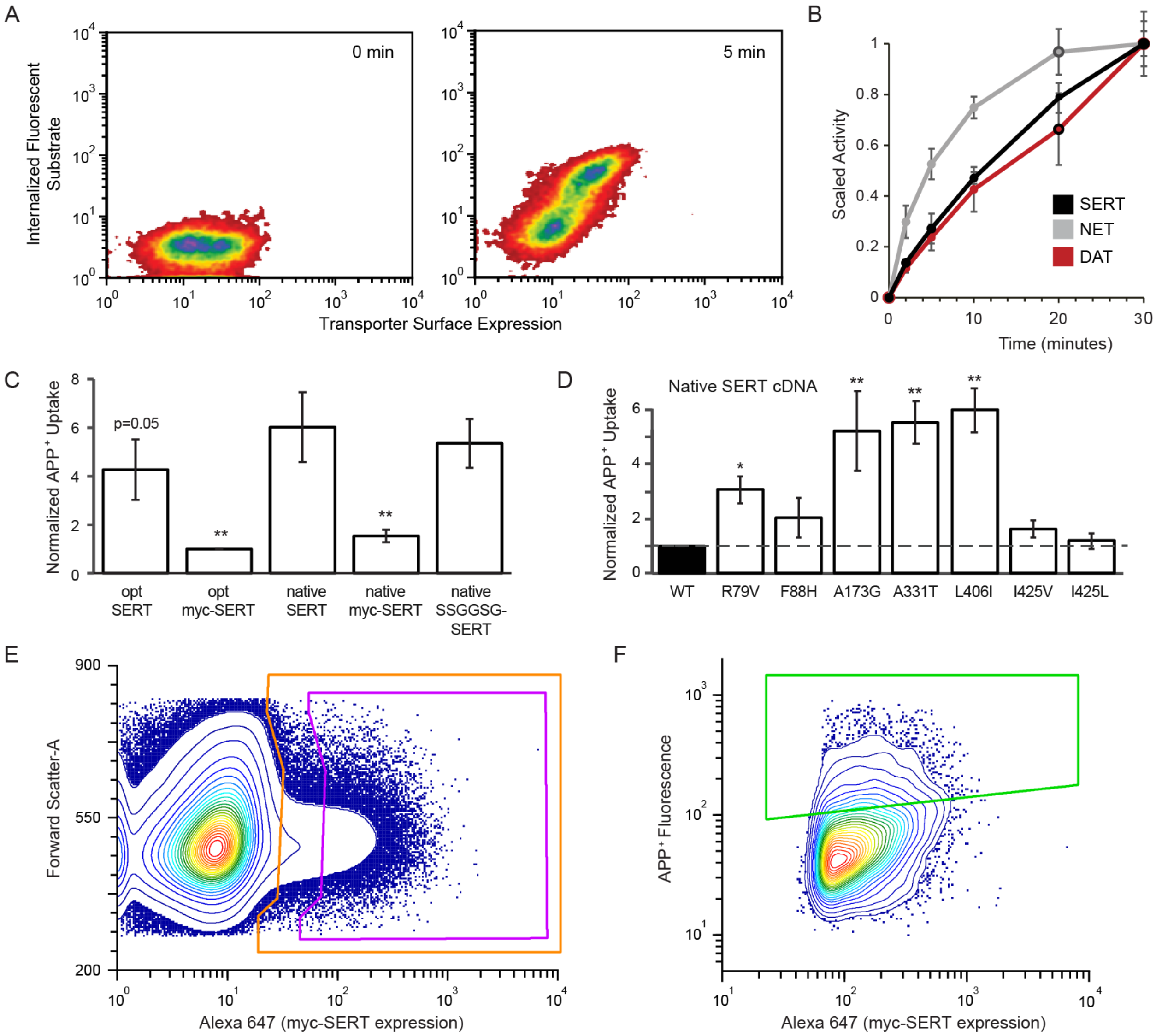

**Figure S2. A flow cytometry-based assay for monoamine neurotransmitter transporter activity.**

**(A)** Transfected Expi293F cells were incubated with 3 μM APP+ and stained with Alexa Fluor 647-anti-myc to detect transporter surface expression.

**(B)** Time course for 3 μM APP+ uptake by Expi293F cells expressing myc-tagged DAT, NET or SERT. The c-myc tags were inserted in ECL2 between Asp200-Ser201, Gly197-Asn198, and Asp216-Asn217, respectively. Nonspecific uptake by untransfected cells is subtracted. ($n = 3$, mean ± SD)

**(C)** In this study, human codon-optimized SERT with a c-myc tag was mutationally scanned; a synthetic codon optimized gene removes sequence elements that are intractable to PCR-based mutagenesis and assists library construction, while the c-myc tag enables detection of surface expressed protein. Activity is comparable between cells expressing SERT from the codon-optimized gene (opt SERT) versus native cDNA (native SERT). The presence of the c-myc tag decreases activity. This is due to the c-myc tag sequence and/or length rather than the choice of insertion site, since a 6-residue spacer (SSGGSG) between Asp216-Asn217 has no effect.

Transfected cells were incubated with 0.5 μM APP+ for 10 minutes. * $p < 0.05$, ** $p < 0.01$, ordinary one-way ANOVA with Dunnett test, constructs/mutants are compared to native SERT. ($n = 3$-7, mean ± SD)

**(D)** Predicted GOF mutations for APP+ import were introduced into the SERT native cDNA without any epitope tag and transiently expressed. Cells were incubated for 2 minutes with 0.5 μM APP+. Wild type SERT is black. * $p < 0.05$, ** $p < 0.01$, ordinary one-way ANOVA with Dunnett test, constructs/mutants are compared to WT SERT. ($n = 7$, mean ± SD)

**(E)** Cells transfected with codon-optimized myc-SERT SSM libraries were incubated with 0.5 μM APP+ and stained with Alexa Fluor 647-anti-myc. Under the transfection conditions, cells typically acquire no more than one sequence variant and most of the population is negative. The top 50% of expressing cells (purple gate) were collected from the SERT-positive population (orange gate).

**(F)** When sorting for APP+ import, the top expressing cells were further gated for the 15% with highest APP+ fluorescence (green gate).

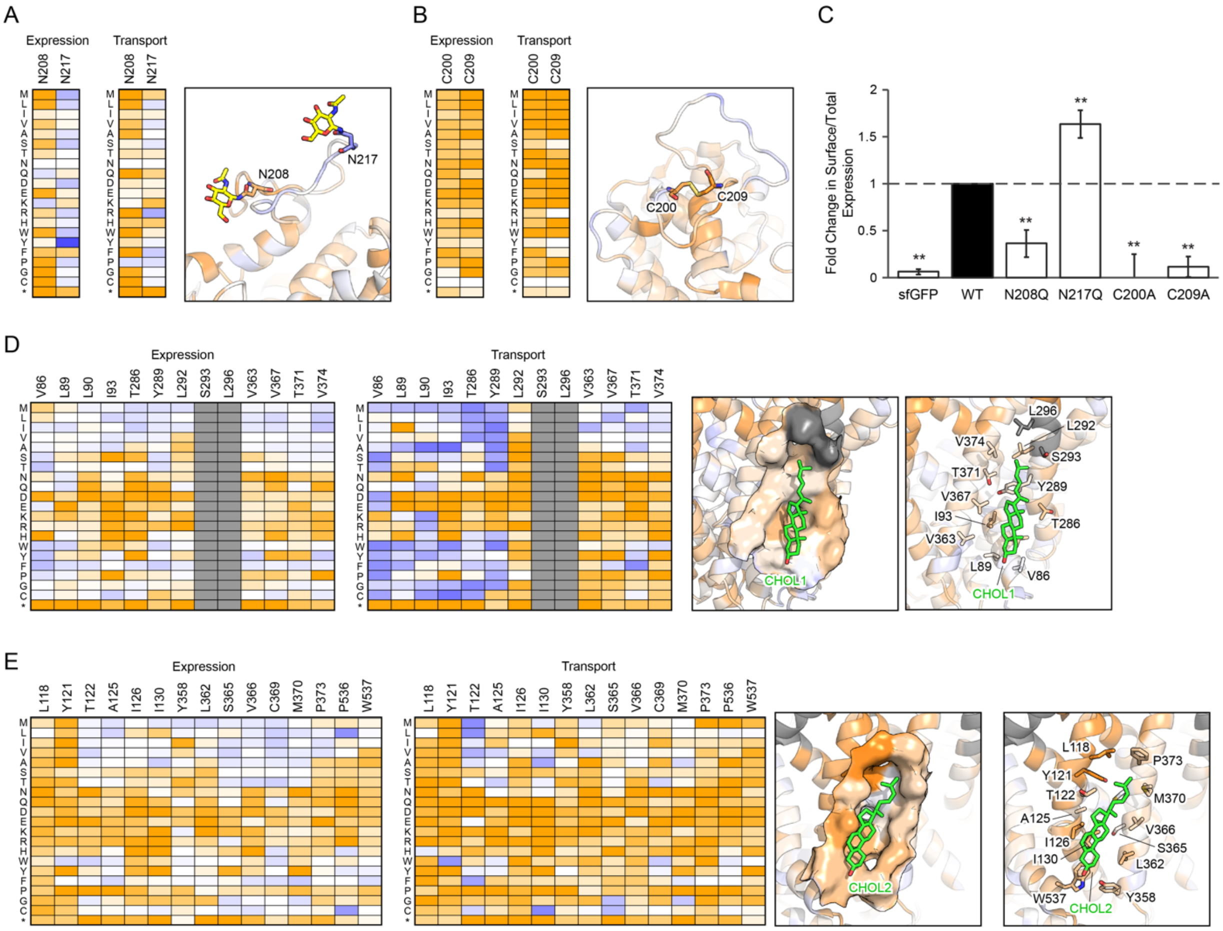

**Figure S3**: **The deep mutational scan identifies residues important for SERT trafficking.** (**A-B**) Heat maps of mutation effects for SERT surface expression and APP+ import for the (**A**) N-linked glycosylation sites and (**B**) the disulfide bond between Cys200 and Cys209. $Log_2$ enrichment ratios are plotted from ≤ -3 (depleted mutations, orange) to 0 (neutral, white) to ≥ +4 (enriched, dark blue). Accompanying snapshots show the SERT structure (PDB: 5I73) as a cartoon representation with residues colored according to conservation score for expression.
(**C**) SERT surface expression based on flow cytometry detection of the c-myc tag normalized by total expression as measured by fluorescence of GFP fused to the N-terminus of SERT. * $p < 0.05$, ** $p < 0.01$, ordinary one-way ANOVA with Dunnett test, constructs/mutants are compared to wildtype SERT. (n = 4, mean ± SD)
(**D-E)** Heat maps of SERT surface expression and APP+ import for two putative cholesterol sites, CHOL1 formed between TM1a, TM5, and TM7 (**D**) and CHOL2 formed between TM2 and TM7. Accompanying snapshot show the SERT structure (PDB: 5I73) as a surface and cartoon representation with residues colored according to conservation score for expression. Cholesterol molecules are placed based on the dopamine transporter (PDB: 4XP1). Mutations with less than 10 reads in the naïve libraries are colored in grey.

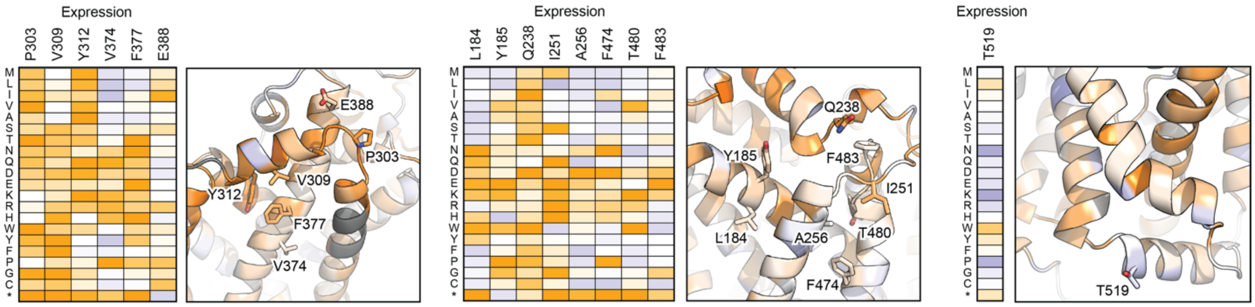

**Figure S4**: **Effects of mutations for SERT surface expression for the top 15 individually co-evolved residues.** $Log_2$ enrichment ratios are plotted from ≤ -3 (depleted mutations, orange) to 0 (neutral, white) to ≥ +4 (enriched, dark blue). Accompanying snapshots show the SERT structure (PDB: 5I73) as a cartoon representation with residues colored according to conservation score for expression. Co-evolutionary strength was calculated using the EVcouplings package.

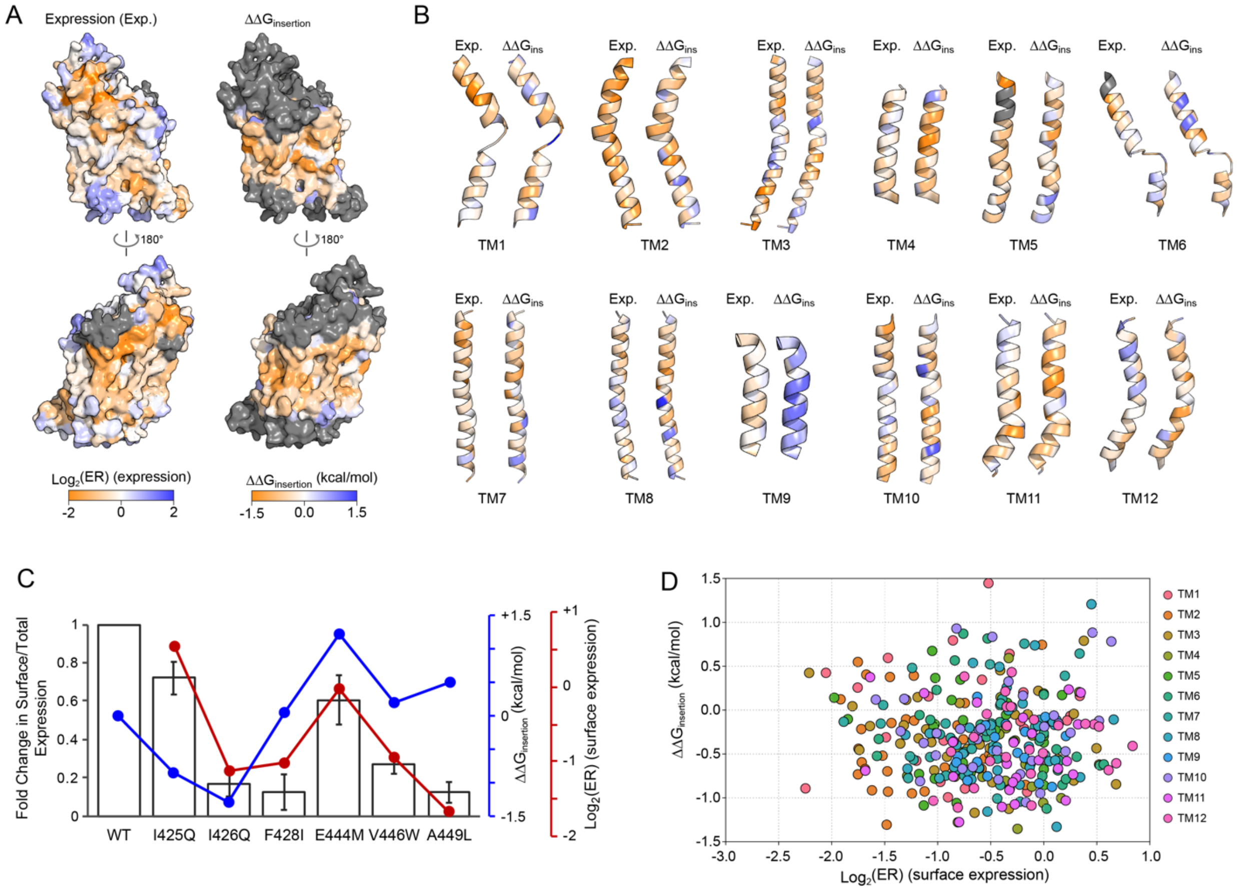

**Figure S5**: **The deep mutational scan for SERT surface expression does not correlate with membrane helix insertion energetics.**

**(A)** Comparison of mutation enrichment ratios for surface expression (left) and predicted ΔΔG of helix membrane insertion (right). The conservation scores for surface expression for each residue are projected on the left three-dimensional surface of SERT and colored from intolerant of mutations (orange) to mutations are enriched (blue). Likewise, the average $\Delta\Delta G_{insertion}$ for each residue is projected on the right three-dimensional surface of SERT. The $\Delta\Delta G_{insertion}$ for an individual residue was averaged for each amino acid substitution and calculated by the difference between ΔGmut and ΔGwild-type. Predicted ΔG values were obtained from the ΔG prediction server v1.0 (*134*).

(**B**) Conservation scores for surface expression and predicted $\Delta\Delta G_{insertion}$ projected on individual transmembrane helices of SERT. Color scheme is that of panel A.

(**C**) SERT surface expression based on detection of the c-myc tag normalized by total expression as measured by fluorescence of GFP fused to the N-terminus of SERT. Five mutations on TM8 predicted by $\Delta\Delta G_{insertion}$ calculations (blue line) to decrease (negative values) or increase (positive values) membrane helix insertion were evaluated. Experimental measurements of surface expression (black bars) instead correlated more closely with enrichment ratios from the deep mutational scan (red line), suggesting proper SERT expression and trafficking is not solely dependent on membrane helix biogenesis.

(**D**) Surface expression conservation scores for each residue are plotted against predicted $\Delta\Delta G_{insertion}$. Individual helices are colored accordingly. $R^2 = 0.000344$, Spearman's $\rho = 0.0083$).

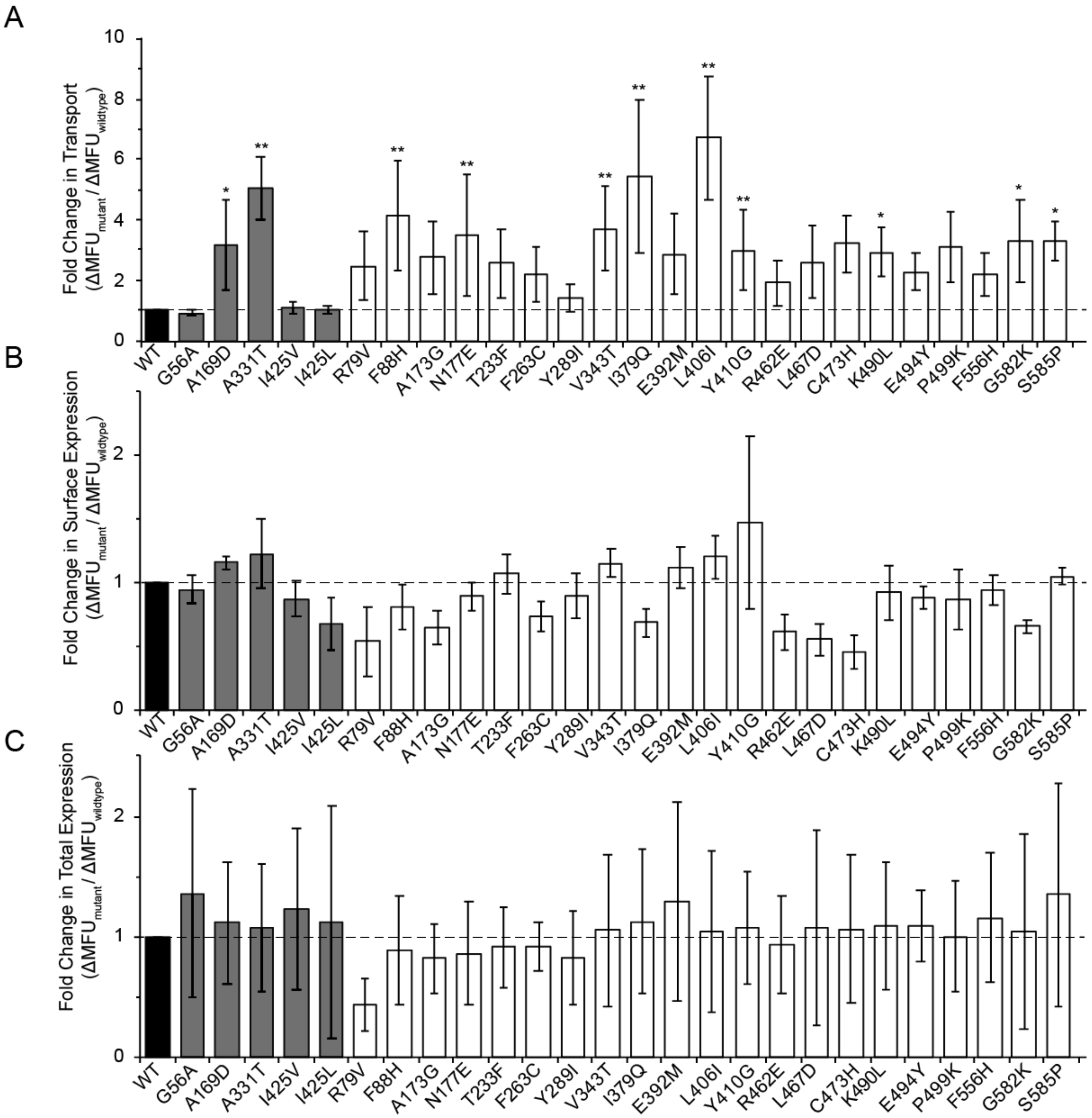

**Figure S6. Validation of gain-of-function (GOF) mutations for APP+ import predicted from the deep mutational scan.**

**(A)** APP+ uptake was measured in Expi293F cells expressing the indicated myc-SERT mutants. Wild type SERT is black. Coding variants reported in databases of human variation are grey. A selection of 22 predicted GOF mutations from the deep mutational scan (white, plus A169D was independently selected as part of this set) were chosen to broadly cover the entire length of SERT. These data show transport activity without any adjustment for differences in SERT surface expression. (n = 4-5, mean ± SD)

**(B)** Surface expression based on detection of the c-myc tag by flow cytometry. (n = 4-7, mean ± SD)

**(C)** Total SERT expression was measured after fixation and partial membrane solubilization. (n = 7-8, mean ± SD)
In all panels, * $p < 0.05$, ** $p < 0.01$, ordinary one-way ANOVA with Dunnett test for comparison to WT.

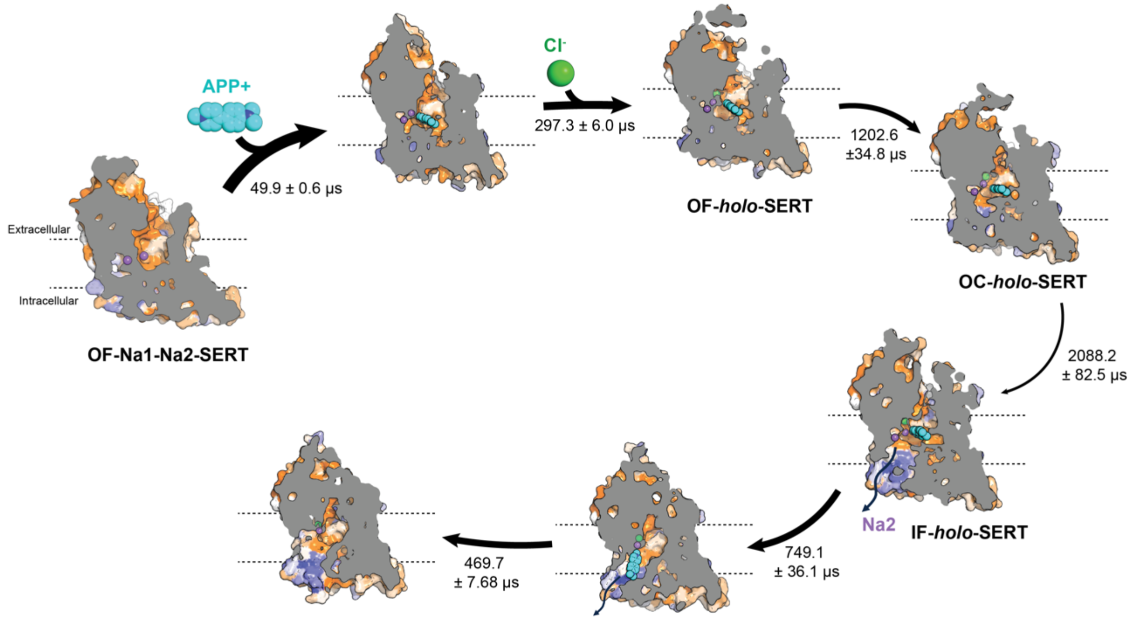

**Figure S7. Major flux pathway for APP+ import.**

The major flux pathway for binding events and conformational changes associated with $Na^+$/APP+ symport initiated from an OF $Na^+$-bound SERT structure. A cross-section through SERT is shown to reveal opening or closing of cavities, colored by conservation score from deep mutagenesis for APP+ import (orange, conserved; dark blue, residues where gain-of-function mutations are enriched). Arrow thickness represents the estimated mean first passage time determined by the Markov state model. Standard deviation is measured by a Bayesian Markov estimator using 100 samples. We note that the rates may not be fully representative of *in vivo* conditions as the simulations are unable to model the cytoplasmic terminal domains as they remain structurally unresolved.

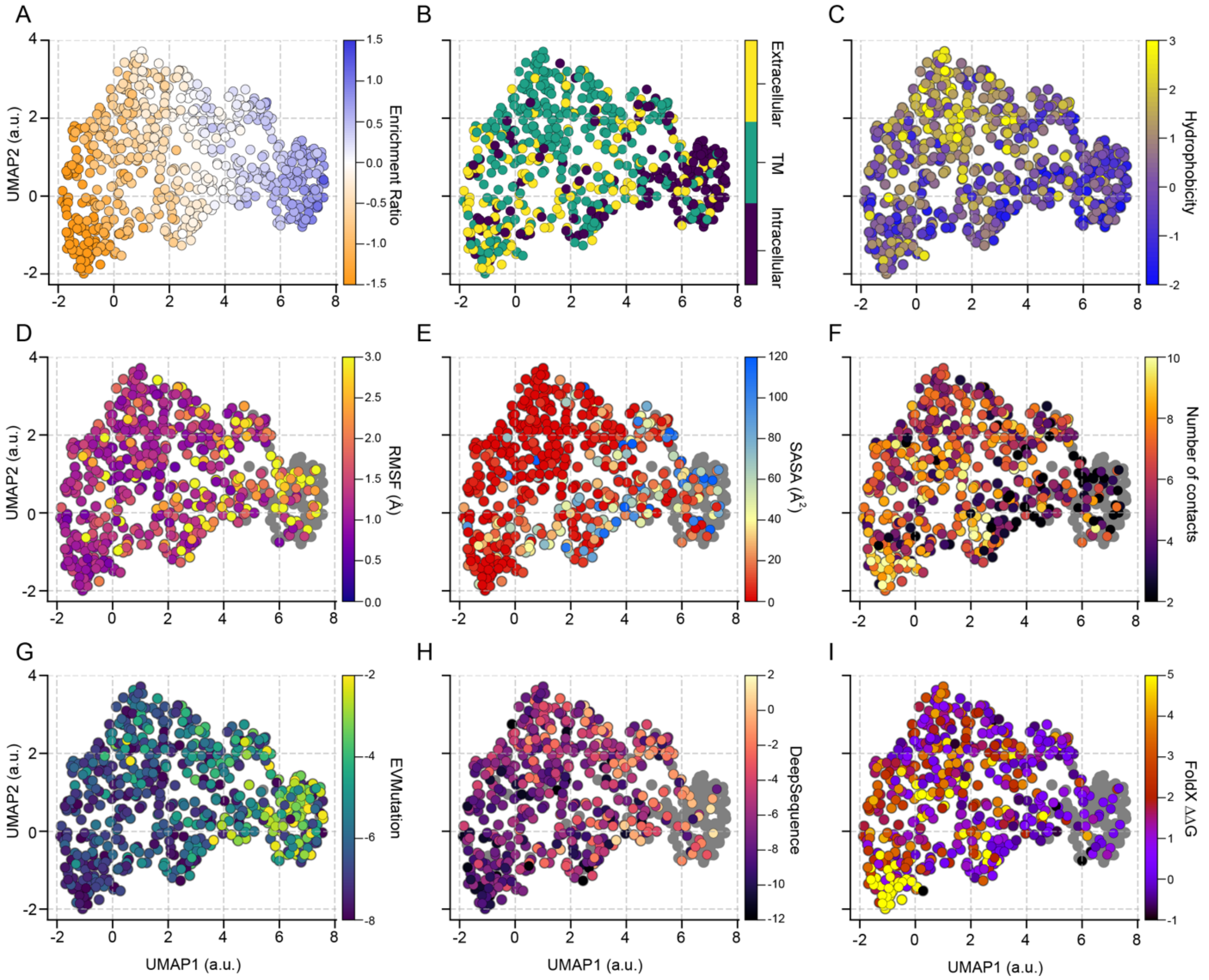

**Figure S8. UMAP projection of the mutational landscape for SERT surface expression.** Each residue was clustered using the UMAP learning algorithm based on its activity following all possible amino acid substitutions. Residues are colored based on different biochemical and biophysical properties: (**A**) surface expression enrichment ratio, (**B**) residue location, (**C**) Kyte-Doolittle hydrophobicity, (**D**) root mean square fluctuation (RMSF) (**E**) solvent accessible surface area (SASA), and (**F**) average number of contacts. The mutational landscape was also compared to scores from variant effect predictors (**F**) EVmutation, (**G**) DeepSequence, and (**I**) FoldX. Residues in which data is unavailable are colored gray. The first UMAP dimension strongly correlates with enrichment ratio (expression $\rho = 0.960$).

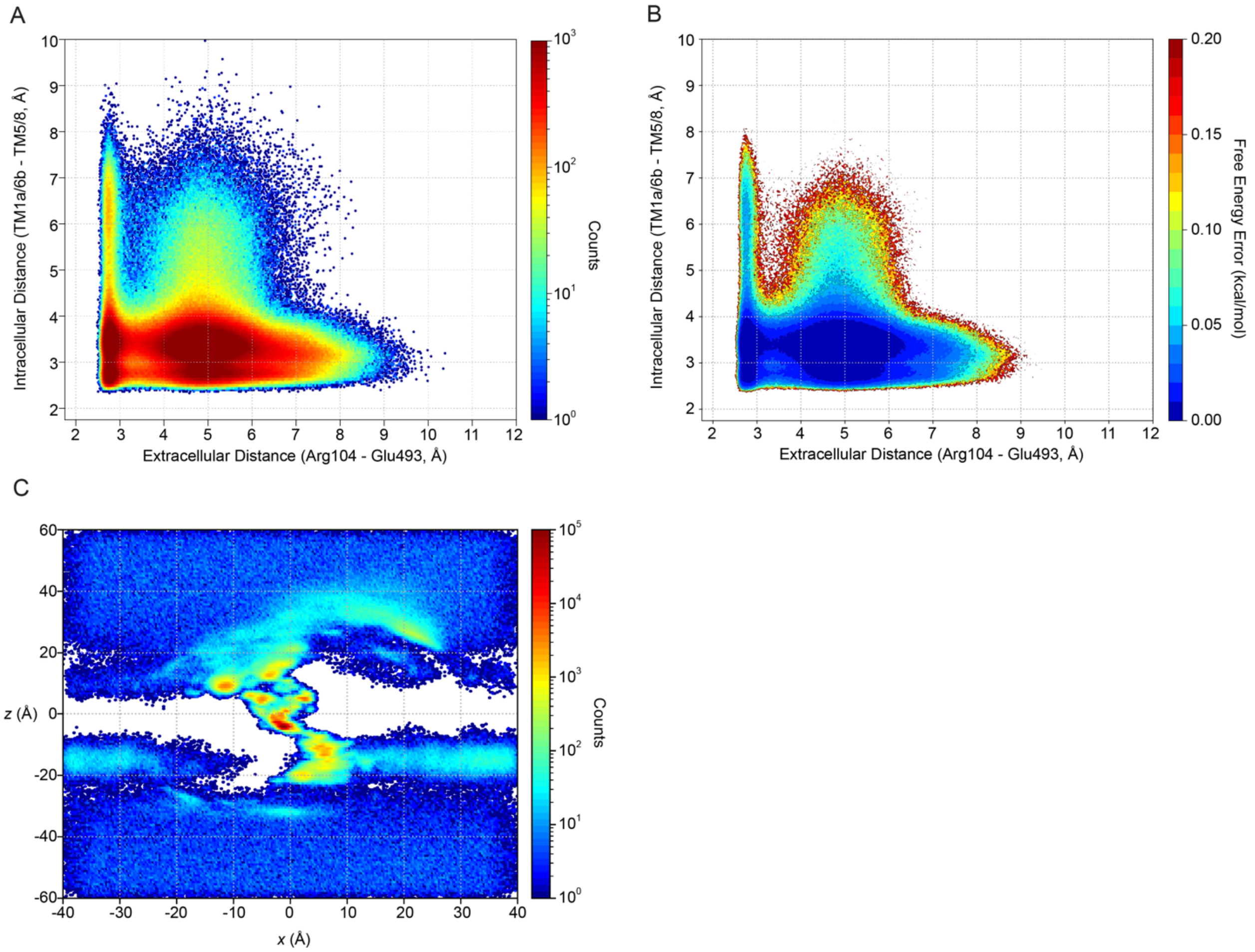

**Figure S9.** (A) Two-dimensional histogram of the simulation data projected on the metrics of extracellular and intracellular distances. (B) Standard free energy error of the conformational landscape. Calculated error of the free energy landscape for SERT-catalyzed APP+ import (APP+-SERT) projected on the metrics of extracellular and intracellular distances. (C) Two-dimensional histogram of the APP+ *x* and *z* coordinate of the simulation box. Negative *z* indicates the intracellular side of the membrane, while positive *z* is the extracellular side.

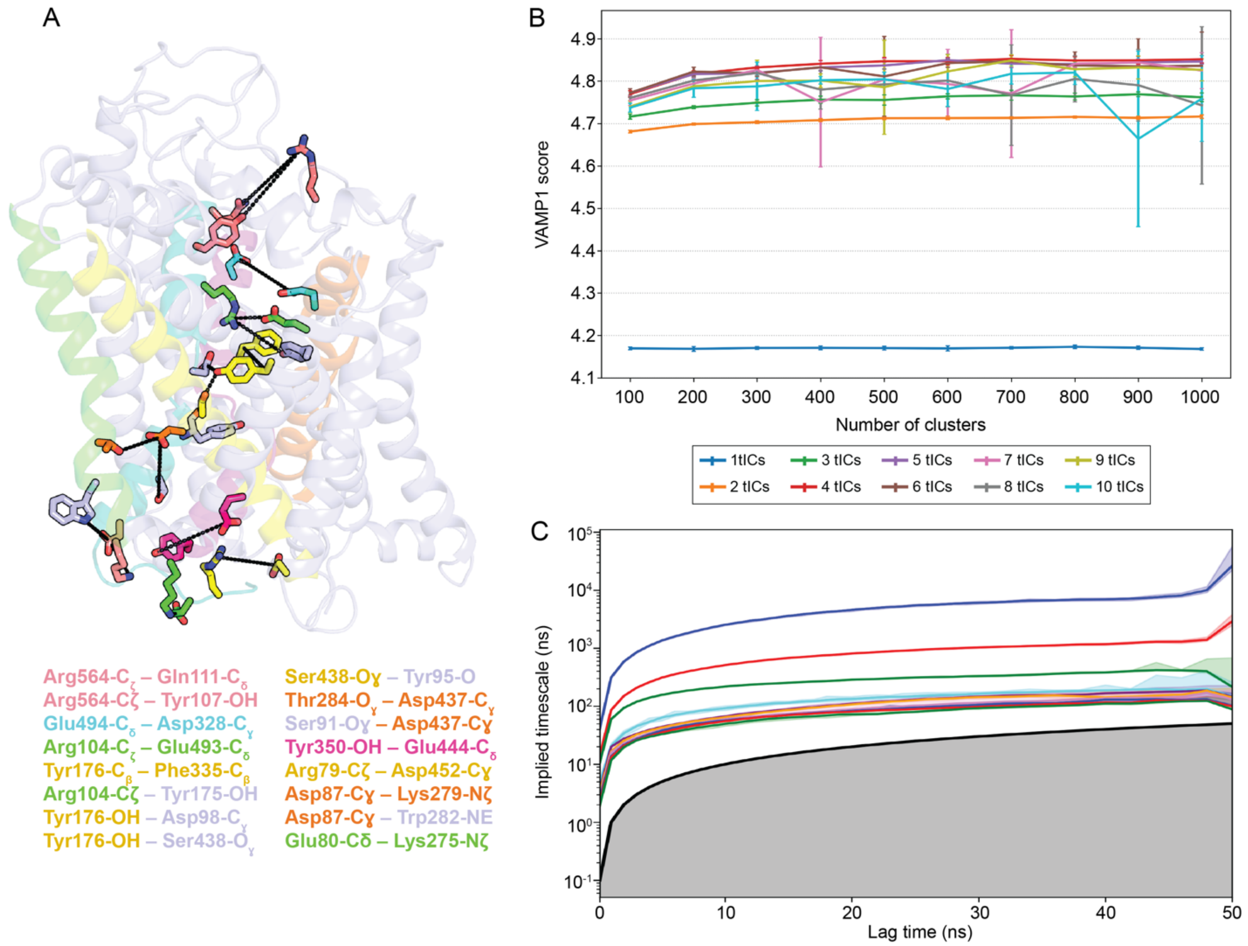

**Figure S10. Feature selection and parameters for MSM construction.**

**(A)** 16 interhelical distances (residue pairs listed in color) along the permeation pathway, together with the z-component of APP+, $Cl^-$, and the symported $Na^+$, were used as featurization metrics for MSM construction. OF-SERT is shown as a pale blue ribbon. TM 1, 5, 6, 8, and 10 are highlighted in teal, green, magenta, yellow, and orange, respectively.

**(B)** Clustering size and time-independent components (tICs) optimization by maximizing the VAMP1 score. As a result, 600 clusters and 4 tICs were chosen to construct the MSM.

**(C)** Implied timescales plots from the transition probability matrix for the first 10 eigenvalues of each MSM constructed using 600 clusters and 4 tICs. Implied timescale curves level off at ~12 ns lag time suggesting Markovian nature. Final MSM was constructed at lag time of 12 ns.

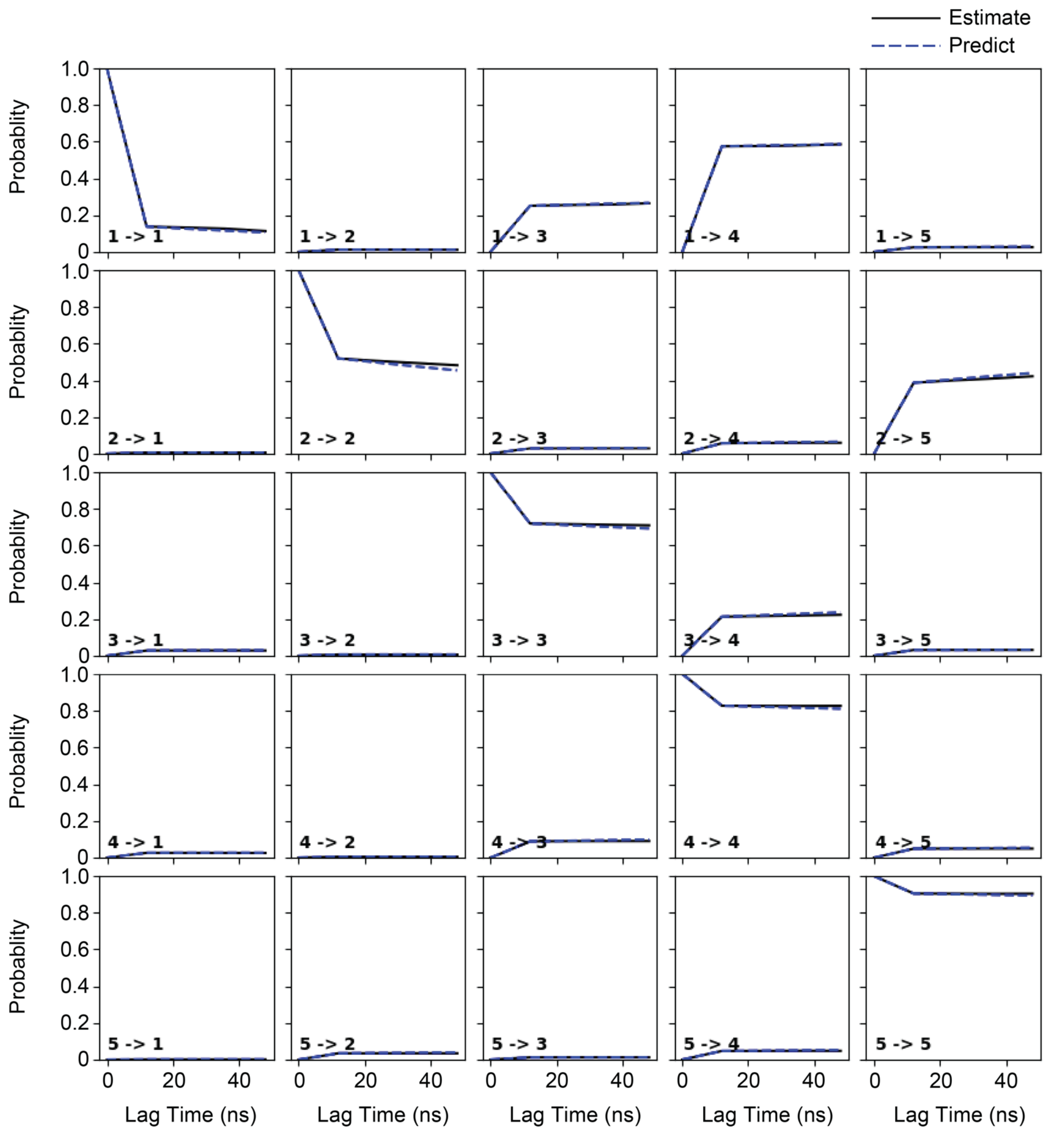

**Figure S11. MSM validation using the Chapman-Kolmogorov test.**
The Chapman-Kolmogorov test evaluates the constructed MSM by predicting the transition probability matrix at greater lag times. Probability of transitions as a function of lag time for a 5 macrostate MSM was obtained for the original MD trajectory (black solid line) and propagation of the MSM (dashed blue line).

**Table S1**: **Conditional probability of coupled events involved in the substrate release process.** The conditional probabilities of the likelihood for either 5-HT or APP+ to exit the orthosteric pocket given certain events during the conformational transport cycle. The metrics described are defined as: the intracellular exit pathway is opened when the minimum distances of the collective intracellular residues is greater than 6Å, Na2 unbound when the distance of the symported $Na^{+}$ ion and Asp452-Cγ is greater than 4Å, Tyr95 flipped when the dihedral angle $\chi 2$ is between -145° and -30°, TM5 unwinding when the helical content of residues 273-282 is less than 0.6, 5-HT exiting the orthosteric pocket when the z-component of center-of-mass of 5-HT is below the z-component of Tyr95-Cα. Error is calculated by estimating the conditional probabilities using 80% of the trajectory data, randomly sampled, to construct the MSM for 500 independent samples. The probability of 5-HT to exit previously computed in (*44*) is listed for comparison.

| | **Probability of APP+ exiting the orthosteric pocket** |
|---|---|
| given Na2 is unbound | 0.202 ± 0.005 |
| given Tyr95 is flipped | 0.096 ± 0.003 |
| given TM5 unfolds | 0.117 ± 0.005 |
| given the intracellular pathway is opened | 0.591 ± 0.018 |
| given all | 0.763 ± 0.017 |
| | **Probability of 5-HT exiting the orthosteric pocket** |
| given Na2 is unbound | 0.277 ± 0.005 |
| given Tyr95 is flipped | 0.562 ± 0.030 |
| given TM5 unfolds | 0.023 ± 0.001 |
| given the intracellular pathway is opened | 0.394 ± 0.007 |
| given all | 0.811 ± 0.068 |

**Table S2: Correlation between biophysical properties and variant effect predictors with UMAP dimensions and DMS enrichment ratio.** Values presented are Spearman's ρ.

| | Expression | | | APP+ Transport | | |
|---|---|---|---|---|---|---|
| **Property** | **UMAP1 Dim.** | **UMAP2 Dim.** | **DMS Enrich. Ratio** | **UMAP1 Dim.** | **UMAP2 Dim.** | **DMS Enrich. Ratio** |
| DMS expression | 0.960 | 0.202 | 1.00 | N/A | N/A | N/A |
| DMS transport | N/A | N/A | N/A | 0.966 | -0.013 | 1.00 |
| Hydrophobicity | -0.249 | 0.358 | -0.184 | -0.130 | -0.408 | -0.098 |
| RMSF | 0.298 | -0.032 | 0.294 | 0.143 | 0.191 | 0.161 |
| SASA | 0.423 | -0.166 | 0.362 | 0.150 | 0.332 | 0.165 |
| No.of contacts | -0.301 | -0.078 | -0.329 | -0.249 | -0.024 | -0.258 |
| EVcouplings | 0.486 | 0.116 | 0.506 | 0.345 | 0.085 | 0.338 |
| DeepSequence | 0.382 | 0.213 | 0.409 | 0.281 | -0.095 | 0.275 |
| FoldX ΔΔG | -0.385 | -0.138 | -0.406 | -0.272 | -0.074 | -0.288 |
